# Comprehensive phylogenetic analysis of the ribonucleotide reductase family reveals an ancestral clade and the role of insertions and extensions in diversification

**DOI:** 10.1101/2022.04.23.489257

**Authors:** Audrey A. Burnim, Matthew A. Spence, Da Xu, Colin Jackson, Nozomi Ando

**Author notes:** These authors contributed equally to this work (Audrey A. Burnim, Matthew A. Spence, Da Xu).

## Abstract

Ribonucleotide reductases (RNRs) are used by all organisms and many viruses to catalyze an essential step in the *de novo* biosynthesis of DNA precursors. RNRs are remarkably diverse by primary sequence and cofactor requirement, while sharing a conserved fold and radical-based mechanism for nucleotide reduction. Here, we structurally aligned the diverse RNR family by the conserved catalytic barrel to reconstruct the first large-scale phylogeny consisting of 6,779 sequences that unites all extant classes of the RNR family and performed evo-velocity analysis to independently validate our evolutionary model. With a robust phylogeny in-hand, we uncovered a novel, phylogenetically distinct clade that is placed as ancestral to the classes I and II RNRs, which we have termed clade Ø. We employed small-angle X-ray scattering (SAXS), cryogenic-electron microscopy (cryo-EM), and AlphaFold2 to investigate a member of this clade from *Synechococcus* phage S-CBP4 and report the most minimal RNR architecture to-date. Using the catalytic barrel as a starting point for diversification, we traced the evolutionarily relatedness of insertions and extensions that confer the diversity observed in the RNR family. Based on our analyses, we propose an evolutionary model of diversification in the RNR family and delineate how our phylogeny can be used as a roadmap for targeted future study.

## Introduction

The wealth of genomic and metagenomic sequence data that has exploded in recent years provides a new opportunity to reexamine enzyme families using large-scale and robust bioinformatic analyses. A particularly important enzyme family that necessitates such analyses are the ribonucleotide reductases (RNRs). RNRs are used by all organisms and many viruses for the conversion of ribonucleotides to 2ʹ-deoxyribonucleotides in the *de novo* biosynthesis of DNA precursors (Torrents et al., 2002). RNRs are especially fascinating from an evolutionary perspective as they exhibit high diversity in primary sequence and cofactor requirement (Lundin et al., 2015), yet they share a common fold and radical-based catalytic mechanism for nucleotide reduction (Licht et al., 1996).

Based on current biochemical evidence, the full catalytic cycle is thought to involve three steps in all RNRs: cofactor-mediated generation of a thiyl radical at a conserved cysteine in the active site, nucleotide reduction, and reduction of active-site residues that provide reducing equivalents for nucleotide reduction (Greene et al., 2020; Holmgren and Sengupta, 2010). Despite being a diverse enzyme family, the core structure of the catalytic subunit (known as α) is a conserved 10-stranded α/β barrel with the thiyl radical on the so-called “finger loop” which connects the two halves of the barrel (**Figure 1**) (Uhlin and Eklund, 1996). RNRs have been biochemically classified into three major groups based on the cofactor used to generate the thiyl radical. Class I RNRs use a ferritin subunit (β) to house a stable radical cofactor and are further subclassified by the metal content of the cofactor (Cotruvo and Stubbe, 2011; Ruskoski and Boal, 2021). For every turnover, the α and β subunits must form a complex to engage in long-range radical transfer to the active site (Kang et al., 2020; Seyedsayamdost et al., 2007). In class II RNRs, the thiyl radical is generated by a 5ʹ-deoxyadenosyl radical (5ʹ-dAdo•) produced from an adenosylcobalamin (AdoCbl) cofactor bound within the active site, and thus these enzymes do not require additional subunits (Licht et al., 1996). In class III RNRs, the thiyl radical is generated by a glycyl radical on the C-terminal domain of the α subunit, which itself is generated by a separate activase enzyme that utilizes *S*-adenosylmethionine (AdoMet) bound to a [4Fe-4S] cluster for radical chemistry (Wei et al., 2014b). Once the thiyl radical is produced in the active site, nucleotide reduction proceeds via a conserved mechanism for all classes (Licht et al., 1996).

**Figure 1.**
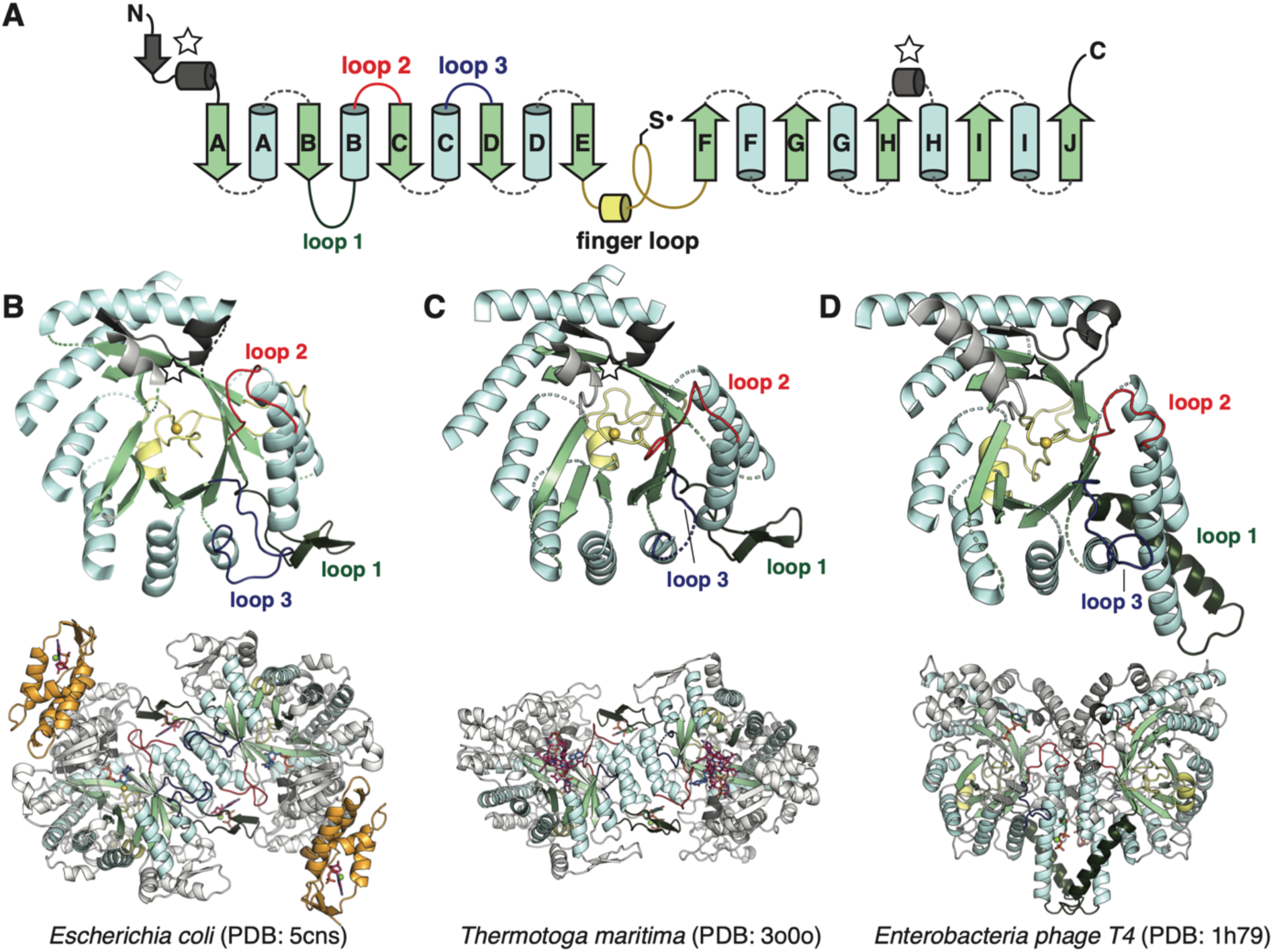
The catalytic fold of the ribonucleotide reductase (RNR) family is a unique 10-stranded ɑ/β barrel, consisting of ten β-strands (light green) and eight ɑ-helices (light blue). (**A**) Each half of the barrel contains a 5-stranded parallel β-sheet (βA-βE and βF-βJ) that is arranged in anti-parallel orientation with respect to each other. The two halves are connected by the so-called “finger loop” (yellow) which typically begins with a short ɑ-helix and contains a conserved cysteine that has been shown to be the site of the catalytically essential thiyl radical in all biochemically characterized RNRs. Diversity among the RNRs is generated by N- and C-terminal extensions as well as the insertions (dashed curves) between the secondary structure elements in the ɑ/β barrel. Loops 1-3 (dark green, red, blue) have special names in the RNR literature for their involvement in specificity regulation. The gray secondary structure elements (starred) are partially integrated in the ɑ/β barrel and are involved in substrate binding. (**B**) The barrel portion (top) of the *E. coli* class Ia catalytic subunit (bottom dimer). The ATP-cone domain is colored in orange. (**C**) The barrel portion (top) of the *T. maritima* class II catalytic subunit (bottom dimer). (**D**) The barrel portion (top) of the T4 phage class III catalytic subunit (bottom dimer). In class III RNRs, the loop 1 region (dark green) is a long helix that is involved in dimerization.

Extensions and insertions about the conserved 10-stranded α/β barrel provide RNRs with mechanisms of allosteric regulation and active-site reduction. Of these, mechanisms to regulate substrate specificity are most integral to the barrel structure. All extant RNRs, with the exception of the monomeric subset of class II RNRs, are thought to dimerize with the αA and αB helices from each monomer forming a four-helical bundle at the interface. In class I and II enzymes, these helices dimerize in anti-parallel fashion such that each end is capped by a functional insertion called loop 1 (**Figure 1B-C**, dark green), which serves as the binding site of specificity-regulating effectors (Larsson et al., 2004; Zimanyi et al., 2016). This so-called specificity site is allosterically coupled to the active site via another functional insertion called loop 2 (**Figure 1B-C**, red) (Zimanyi et al., 2016), such that the proper balance of nucleotides for DNA synthesis is maintained (Bester et al., 2011; Reichard, 1988). Remarkably, in monomeric class II RNRs, the loop 1 insertion is long enough to interact with the αA and αB helices and structurally mimic the dimer interface and specificity site seen in dimeric class II RNRs (Sintchak et al., 2002). In class III enzymes, the insertion equivalent to loop 1 is a helix (Logan et al., 1999) (**Figure 1D**, dark green), which stabilizes an α2 dimer with the αA and αB helices of the two monomers in parallel orientation, and specificity effectors bind in the dimer interface (Larsson et al., 2001). Compared with specificity regulation, which appears to be important in all RNRs, overall activity regulation is only associated with RNRs that have an N-terminal extension with an ATP-cone motif (**Figure 1B**, orange domain) (Aravind et al., 2000; Jonna et al., 2015; Meisburger et al., 2017) or recently found truncated versions of this motif that serve as a form of convergent allostery (Parker, 2017; Thomas et al., 2019). The C-terminus, on the other hand, is used for enzyme turnover. In class III RNRs, the glycyl radical domain is located at the C-terminus, whereas class I and certain class II RNRs are known to have disordered C-terminal tails with a pair of cysteines that are used for reducing the active site (Booker et al., 1994; Mao et al., 1992). However, in general, much remains unknown about the functional diversity of RNR C-termini.

The ancestor of modern RNRs, and its subsequent evolution into the three known classes has long been of interest for its hypothesized importance in transitioning life from an RNA/protein world to a DNA world (Lundin et al., 2015, 2009; Poole et al., 2002; Reichard, 1993; Stubbe, 2000; Stubbe et al., 2001; Torrents, 2014; Torrents et al., 2002). Class I RNRs have been proposed to be the most recently evolved class as they require molecular oxygen, which became abundant as the Earth’s atmosphere transitioned from anoxic to oxic. In contrast, class III RNRs are often thought to be the most ancient class as glycyl radicals and [4Fe4S] clusters are both extremely oxygen-sensitive (Fontecave et al., 1989) and Fe-S chemistry is thought to have a prebiotic origin (Goldman and Kaçar, 2021). The AdoCbl chemistry used by class II RNRs, on the other hand, is neither oxygen-dependent nor especially oxygen sensitive. Based on the relative oxygen sensitivities and biosynthetic complexities of the different cofactors, class III RNRs have been proposed to be the most likely progenitor of the RNR family (Reichard, 1997; Riera et al., 1997). However, whether class III predates class II has been debated when considering cofactor availability (Stubbe, 2000). Indeed, cobalamin, AdoMet, and Fe-S clusters have all been proposed to be part of the cofactor set used by the last universal common ancestor (LUCA) (Weiss et al., 2016). In an alternate model, dimeric class II and III RNRs evolved in parallel from a shared ancestor using class II-like chemistry, and class I RNRs then evolved from class II (Lundin et al., 2015). Despite these efforts to understand the diversity and evolution of RNRs, the reconstruction of a large-scale, unifying phylogeny that includes all classes of RNRs has been hindered by the significant computational demand imposed by the high sequence diversity of the extant members of the family.

In this study, we used the conserved 10-stranded α/β barrel to anchor highly diverged RNR sequences in a structure-based alignment and used maximum-likelihood inference to reconstruct the largest RNR phylogeny to-date, consisting of 6,779 α sequences, that unifies all classes. Our comprehensive phylogeny shows the parallel development of three major clades, corresponding to the three known classes, with a small, phylogenetically distinct clade, which we denote as class Ø (pronounced “oh”), placed as an ancestral clade to the class I and II RNRs. Using small-angle X-ray scattering (SAXS), cryo-electron microscopy (cryo-EM) and AlphaFold2 (Jumper et al., 2021), we show that the class Ø α is the most minimal RNR structurally characterized thus far. These observations were corroborated by evo-velocity analysis, an alignment-independent method that employs language models to infer sequence fitness and evolutionary trajectories (Hie et al., 2022). Using AlphaFold2 and sequence analysis to characterize insertions and the N- and C-terminal extensions of the RNR family, we further describe mechanisms of RNR diversification in the context of allosteric activity regulation and active-site reduction. Together, our analyses indicate that class III RNRs diverged early on and evolved independently of class I and II RNRs, which diverged from an ancestor shared with the minimal, class Ø RNRs.

## Results

### Phylogenetic reconstruction and evo-velocity analysis reveal the evolution of three RNR classes and a novel clade

To study the molecular evolution of the RNR family, we performed comprehensive phylogenetic inference on the catalytic α subunits (**Figure 2**). Unlike sequence analyses, such as sequence similarity network (SSN) analysis (Atkinson et al., 2009), which provide insight exclusively on the global similarities shared between extant sequences, phylogenetic inference delineates the evolutionary history of related sequences utilizing complex models of evolution that account for the rate of mutation of base pairs or amino acids. However, large-scale phylogenetic reconstruction on entire protein superfamilies is technically challenging and computationally demanding. Homology is often difficult to detect between extensively diverged proteins, complicating the generation of a robust sequence alignment, without which accurate topological reconstruction and model parameterization cannot be achieved. To overcome these challenges, we adapted a recently developed workflow (Spence et al., 2021), which uses ensembles of hidden Markov models (HMMs) guided by structural information to build an accurate alignment of 6,779 α sequences that spans 5 PFAM families.

**Figure 2.**
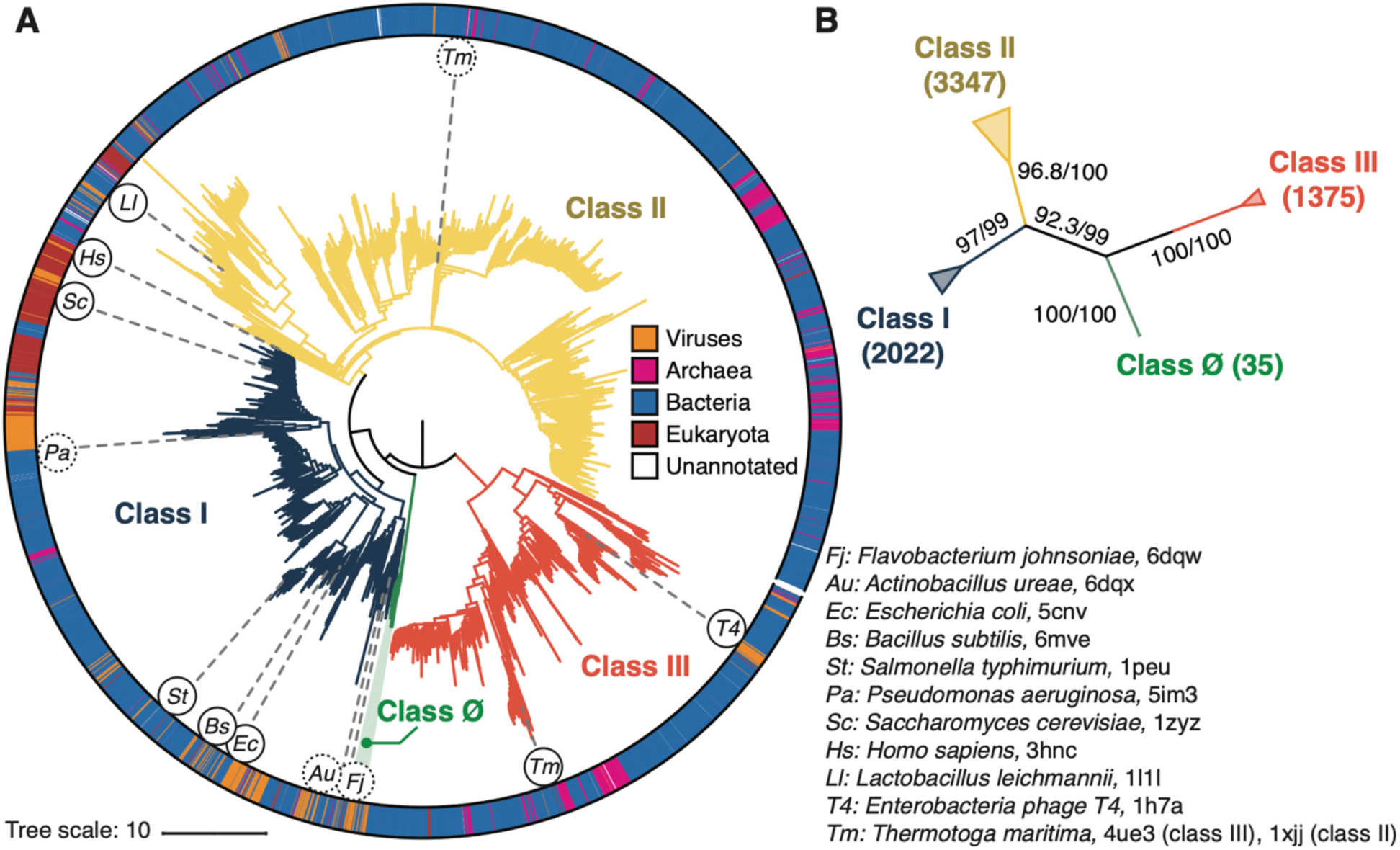
Phylogenetic reconstruction of the RNR superfamily. (**A**) A representative phylogeny of 6779 extant RNR ɑ sequences rooted at the midpoint. The superfamily forms four distinct lineages, with the three major clades (blue, yellow, red) corresponding to the three biochemically known classes I-III. Class III is the most distantly related clade (red). A small ancestral clade (green) forms an outgroup to the clades corresponding to classes I and II (blue and yellow, respectively). The scale bar represents the expected number of amino acid substitutions per site in the alignment. The taxonomic group each sequence belongs to is labelled by a color strip at the circumference. Sequences used for the structural alignment are mapped onto the tree by organism ID in circles. Organism IDs in dashed circles are α sequences that were utilized in the structural alignment but were filtered in favor of more representative sequences for the tree inference and thus are not on the tree (see Methods). These structures are represented by closely related sequences with ≥ 80% sequence identity. (**B**) An unrooted representation of the RNR phylogeny with branch supports (UFboot/SH-ALRT statistics) for the placements of major lineages. Deep nodes in the RNR phylogeny are resolved with high confidence.

Phylogenetic reconstruction was performed by maximum-likelihood (ML) inference. Tree searches were performed with 10 replicates under the sequence evolution model LG+R10 (Kalyaanamoorthy et al., 2017; Le and Gascuel, 2008), which was selected by Akaike and Bayesian information criteria (Dridi and Hadzagic, 2019) on a representative subset of the full 6,779 RNR sequences. To test the robustness of our phylogenetic hypotheses against model diversity, we inferred an additional 10 replicates of tree-search under an alternative general amino acid replacement matrix (WAG+R10) (Whelan and Goldman, 2001). Each of the 10 phylogenies inferred under LG+R10 failed rejection by the approximately unbiased (AU) test (Shimodaira, 2002) conducted to 10,000 replicates (lowest P-value = 0.283), including those whose branch supports failed to converge. All 20 inferences, irrespective of evolutionary model, converged on a similar phylogenetic topology with the same cladistic groups (**Figure 2 – figure supplement 1**), indicating that the likelihood surface of the evolutionary relationships of RNR α subunits has a well-defined global maximum that we had reached in our tree-searches. This was corroborated by high branch supports (ultrafast bootstrap approximation 2.0 and approximate likelihood ratio tests) for major bifurcations in each independent reconstruction, including deep nodes at the midpoint root of each tree (**Figure 2B**). The topology that is presented in **Figure 2** had the highest branch supports at key nodes of interest in RNR evolution.

Consistent with our current understanding of RNR evolution, all reconstructed topologies that converged and failed rejection by the AU-test resolved the three biochemical classes of RNRs as monophyletic lineages (**Figure 2**, blue/yellow/red). Additionally, we find strong support for classes I and II diverging from a single common ancestor that shared an ancestor with the class III RNRs. We identified a novel clade of α sequences that diverged from the last common ancestor (LCA) of classes I and II, which we denote as the class Ø clade (**Figure 2**, green). The branch that separates the LCA of classes Ø, I, II from class III is the midpoint of the phylogeny. Rooting on this branch places class III as the most ancestral, followed by the emergence of the LCA of classes Ø, I, II and the subsequent divergence of classes I and II.

As phylogenetic inference is susceptible to biases introduced by taxon sampling (Pollock et al., 2002), model parameterization and assumptions, sequence alignment (Simmons et al., 2011), and tree-search, we also used a recently described phylogeny-independent method known as evo-velocity analysis to test our phylogenetic hypotheses (Hie et al., 2022). Evo-velocity is a machine-learning analysis based on the principles of natural language processing (NLP) (Hie et al., 2022). NLP models such as ESM-1b, a >650,000,000 parameter NLP model trained on UniProt50 amino acid sequences (Rives et al., 2021), can embed query sequences in a high-dimensional space where likelihoods can be numerically computed as proxy fitness scores on a graph network. By assuming that the inherent directionality of the embedded sequence graph-network moves in the direction of evolution, as evolution compels proteins towards more fit (i.e., more probable in the NLP model) sequences, sequences with low learned probabilities can be interpreted as starting points in evolutionary trajectories. Evo-velocity thus bypasses many of the assumptions of phylogenetic inference as it is alignment independent, does not require model parameterization or treesearch, and has previously been used to validate conclusions based on superfamily-scale phylogenetic inference (Hie et al., 2022; Spence et al., 2021).

When projected onto a two-dimensional basis (see **Methods**), the ESM-1b embedded RNR graph network (**Figure 3**) shares many common topological features with the full family phylogeny (**Figure 2**). Class III RNRs (**Figure 3A**, red) belong to a diverged cluster of sequences that have evolved independently of class I and II sequences (**Figure 3A**, blue/yellow). Conversely, class I and II are joined over the same sequence space, indicating a common recent ancestor. The class Ø sequences occupy the interface between classes I and II (**Figure 3A**, green), consistent with their phylogenetic placement as ancestral to the class I/II lineage. Evo-velocity analysis also identified multiple roots in the RNR family (**Figure 3B**, dark purple). This is likely a consequence of sequences diverging over geological timescales and lacking intermediate phylogenetic information between major groups. The multiple roots belong to each of the major lineages, signifying independent evolution down distinct trajectories. However, pseudotime velocity, a proxy for phylogenetic depth (Haghverdi et al., 2016), clearly captures the interface of class I and II sequences (including class Ø) as the most ancestral within that lineage (**Figure 3C**). The congruence between phylogenetic and NLP analyses supports our hypotheses on the evolution of extant RNRs. The projection of class Ø sequences onto the ancestral interface of classes I and II additionally provides evidence that they indeed hold a pivotal position in RNR evolution.

**Figure 3.**
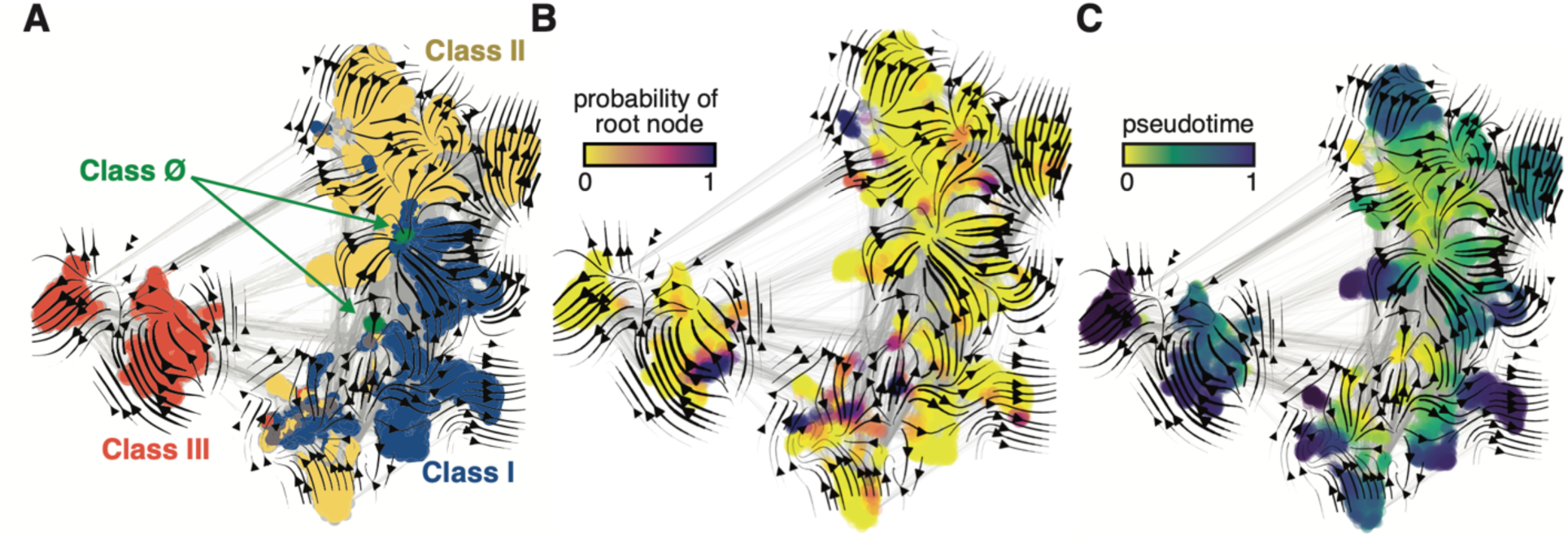
Evo-velocity analysis of the RNR superfamily. ESM-1b embedded RNR sequences were projected onto a two-dimensional vector plot, where the horizontal and vertical axes are UMAP 1 and 2, respectively. Each colored point in the plot corresponds to one of the 6779 ɑ sequences from the full RNR phylogeny. (**A**) Vector field plot colored by RNR classification: clade Ø (green), class I (blue), class II (yellow), class III (red). (**B**) Vector field plot colored by the probability of a sequence being a root of the sequence space, where purple sequences are the most likely to represent probable roots. (**C**) Vector field plot colored by pseudotime, a proxy for phylogenetic depth. Yellow (pseudotime = 0) represents ancestral sequences and indigo (pseudotime = 1) represents sequences that have diverged the most from ancestral sequences.

### Class Ø RNRs are minimal and share features of class I and II enzymes

The class Ø clade consists of RNRs from both bacteria and phages, many of which are phototrophs. Nearly half of the sequences in the clade were obtained from cyanophages (**Figure 4A**, bolded species) or uncultured phages, while the remainder were obtained from marine bacteria, including those collected in the *Tara* Oceans Expedition (Tully et al., 2018). Performing a BLAST search of photosynthetic genes, such as *psbA* and *psbD*, against the genomes (of which many are incomplete) of the 35 bacteria and phages listed in the clade yielded 11 hits (**Figure 4A**, species in green).

**Figure 4.**
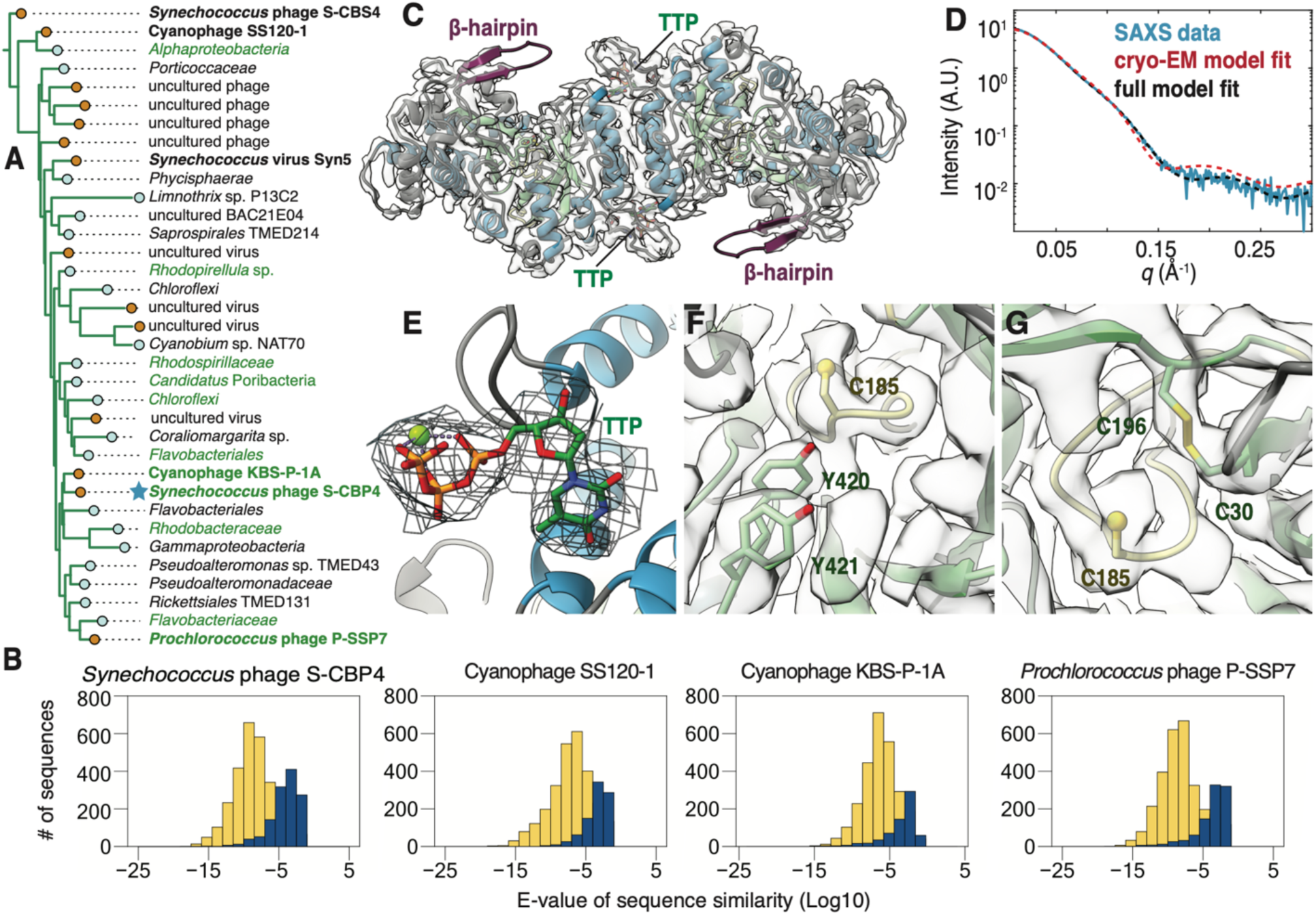
The class Ø α subunit shares similarities with both class I and II α subunits. (**A**) An expansion of the class Ø clade from the full tree in Figure 2. Cyanophage sequences in this clade are bolded. Genomes with identified photosynthetic genes, *psbA* and *psbD*, are colored in green. (**B**) Representative results from an all-vs-all pBLAST search of every class Ø sequence against all RNR α sequences from other clades. Blue bars represent number of significant hits (E-values < 10^-3^) of the title sequence to sequences in the class I clade. Yellow bars represent number of sequences from the class II clade. Overall, the class Ø clade shares greater homology with the class II RNRs. (**C**) A 3.46-Å cryo-EM map of the *Synechococcus* phage S-CBP4 α subunit (shown at a threshold of 2.17) depicts a dimer with TTP bound at the allosteric specificity sites. The β-hairpin (violet) is a shared trait of class I and II RNRs. (**D**) Experimental SAXS profile (blue solid) of the *Synechococcus* phage S-CBP4 α subunit in the presence of 200 μM TTP, 200 mM GDP is explained well by the theoretical scattering of our cryo-EM model (red dashed). Model-data agreement is further improved by modeling the disordered N- and C-termini in AllosModFoXS (black dashed, see Methods) (Schneidman-Duhovny et al., 2010). Cryo-EM density for (**E**) TTP at the specificity site, (F) a stacked tyrosine dyad adjacent to the catalytic cysteine, and (**F**) the oxidized cysteine pair in the active site is shown at a threshold of 2.77.

The classification of the cyanophage sequences observed in this clade (**Figure 4A**, bolded species) has been previously debated (Harrison et al., 2019). These RNRs from *cyanosipho*- and *cyanopodoviruses* were initially annotated as class II, based on studies of the *Prochlorococcus* phage P-SSP7 genome, which contains two consecutive genes that were proposed to represent a split RNR with homology to class II RNRs (Dwivedi et al., 2013; Lindell et al., 2005; Sullivan et al., 2005). More recently, it was discovered that these two genes encode for an RNR ɑ subunit and a small ferritin-like protein and were thus reannotated as class I (Harrison et al., 2019). However, in the phylogenetic analysis reported in this previous study, the cyanophage ɑ sequences fell in either the class I or class II clades depending on the stringency for homology used for grouping the sequences. This result contrasts with our work, where the class Ø clade is phylogenetically distinct from the other major clades (**Figure 2**). Additionally, we find that all completely sequenced operons with a class Ø RNR ɑ gene (24/35 in **Figure 4A**) contain a small ferritin-like gene (annotated with InterPro family number for the ferritin-like superfamily IPR009078) immediately downstream (**Figure 4 – figure supplement 1**), and thus, this feature is not exclusive to cyanophages. Interestingly, an all-vs-all pBLAST search of every class Ø sequence against all class I sequences in our phylogeny detects poor homology (E-value > 0.05) (**Figure 4B**, blue), while querying class Ø RNRs against class II returns hundreds of significant hits (E-value < 10E-10) (**Figure 4B**, yellow). Thus, by sequence homology, class Ø RNRs appear to be more class-II like despite being associated with a ferritin-like gene that is reminiscent of class I RNRs, which supports the phylogenetic results indicting they are a distinct class.

The class Ø lineage consists of α sequences with an average length of 412 amino acids, which are significantly shorter than those of the class Id catalytic subunit that were previously described as having a minimal architecture with a median length of 565 amino acids (Rose et al., 2019). To examine the structural features of the class Ø RNR, we used a combination of AlphaFold2, SAXS, and cryo-EM to characterize the *Synechococcus* phage S-CBP4 α subunit. In the absence of nucleotides, our SAXS profiles are consistent with a monomer-dimer equilibrium (**Figure 4 – figure supplement 2**) where the S-CBP4 α subunit is predominantly monomeric at low protein concentrations (≤4 μM). Addition of substrate (guanosine diphosphate, GDP) does not shift this equilibrium, but the presence of the corresponding specificity effector (thymidine triphosphate, TTP), strongly favors dimerization (**Figure 4 – figure supplement 2**). This result is consistent with TTP binding the specificity site to stabilize the same dimeric form adopted by class I and II α subunits. Using solution conditions that saturate nucleotide binding (200 μM TTP, 200 μM GDP), we obtained a cryo-EM map of the S-CBP4 α2 dimer to 3.46-Å resolution (Fourier shell correlation, FSC = 0.143) (**Figure 4C**, **Figure 4 – figure supplement 3**). An atomic model (residues 22-426 out of 470) was refined against the cryo-EM map using an AlphaFold2 prediction as the starting model (**Table 1**). The elongated shape of the S-CBP4 α2 dimer observed by cryo-EM (**Figure 4D**, red dashed curve) explains the distinct flattened region in our experimental SAXS profile between *q* ∼ 0.05 - 0.15 Å^-1^ (**Figure 4D**, blue curve), and excellent model-data agreement is observed over the full *q*-range when the disordered N- and C-termini are included in the model (**Figure 4D**, black dashed curve and blue curve).

Overall, the S-CBP4 α subunit displays the most minimal RNR architecture discovered to-date with few insertions about the catalytic barrel (**Figure 4 – figure supplement 4**). The minimal set of insertions includes traits shared by class I and II RNRs, such as loop 1, where we find cryo-EM density for bound TTP (**Figure 4E**). Additionally, the S-CBP4 α subunit contains a long β-hairpin following the βH strand of the catalytic barrel that in class II RNRs is thought to be important for cofactor binding (Sintchak et al., 2002) and in class I RNRs is thought to be important for interactions with the β subunit (Kang et al., 2020). Although we do not observe density for the C-terminus past the βJ strand of the catalytic barrel, two consecutive tyrosine residues (Y420 and Y421) on βJ are well-resolved and stacked adjacent to the catalytic cysteine (C185) at the tip of finger loop (**Figure 4F**). In class I RNRs, this stacked arrangement of the tyrosine dyad is required for long-range proton-coupled electron transfer (PCET) between the β subunit and the catalytic cysteine (Greene et al., 2017). By comparison, inspection of existing structures (Larsson et al., 2010; Sintchak et al., 2002) suggests that in class II RNRs, the space between the βJ strand and the finger-loop cysteine forms a pocket to accommodate the adenosyl group of the cobalamin cofactor. Finally, we observe a disulfide between active-site cysteines (C30 and C196) that in class I and II RNRs serve as reducing equivalents during nucleotide reduction (**Figure 4G**). Consistent with an oxidized active site, cryo-EM density for the substrate is not observed.

Although the class Ø ɑ subunit contains a tyrosine dyad poised for PCET like class I ɑ subunits, the class Ø ferritin-like sequences are significantly shorter (average length of 240 amino acids) than those of *bona fide* class I β subunits (average length of 346 amino acids). Querying each complete class Ø ferritin-like sequence returns only a single homologous sequence over the whole length of the alignment (all hits E-value > 1E-4, excluding one) when searched against PFAM families PF00210 (ferritin-like proteins, 17,981 entries) and PF00268 (class I RNR β subunit, 9,898 entries). However, based on AlphaFold2, which detects more than sequence homology, class Ø ferritin-like proteins resemble class I β subunits with shorter insertions and termini (**Figure 4 – figure supplement 5A**-D). As in class I β subunits, the class Ø proteins are predicted to contain a core ferritin fold consisting of two helix-turn-helices (**Figure 4 – figure supplement 5A** and 5C, red/pink and yellow/green), each contributing an E/D + EXXH metal-binding motif (Ruskoski and Boal, 2021). Additionally, while most class I β sequences have a tyrosine serving as the site of the radical downstream of the first metal-binding motif, class Ø ferritin-like proteins contain a redox-inert residue (Phe or Leu) at this site, much like class Ic RNRs (Bollinger et al., 2008; Cotruvo and Stubbe, 2011) (**Figure 4 – figure supplement 5**E). Class Ø ferritin-like sequences also contain a tyrosine (Y239 in **Figure 4 – figure supplement 5A**), which in class I RNRs is involved in inter-subunit radical transfer (Y356 in **Figure 4 – figure supplement 5C**). However, the C-termini of the class Ø ferritin-like proteins are unusually short, terminating at the residue downstream from this tyrosine, and lack the ∼10-residue extension that class I β subunits use for binding to the α subunit. Thus, although the class Ø ferritin-like proteins are likely to use a similar radical-generating mechanism as class I β subunits, they may exhibit key differences in protein-protein interactions involved in PCET. However, ongoing biochemical work to confirm this has thus far been nontrivial, and further investigation will be the focus of future work.

Regardless of biochemical classification, the class Ø α sequences highlight how the consensus catalytic barrel modified with a handful of insertions represents a minimal RNR structure. To understand the mechanisms by which the RNRs may have diversified, we therefore performed detailed analyses of the insertions and extensions about the barrel. These analyses, which are described in the following sections, provide a way to understand the evolutionary model implied by our α phylogeny, namely that class III RNRs diverged early on and that the class I and II RNRs emerged from a common ancestor shared with the minimal class Ø RNRs.

### A single origin for the N-terminal ATP-cone domains

We first investigated mechanisms of RNR diversification by examining the N-terminal extensions. This region exhibits significant diversity in length due to the variable numbers of ATP-cone motifs (Jonna et al., 2015). The canonical ATP-cone is a ∼100-residue globular domain composed of a four-helical bundle with a three-stranded β-sheet cap (**Figure 1B**, orange domain). A functional ATP-cone domain houses an allosteric site that is responsible for regulating overall enzyme activity in response to endogenous ATP and dATP concentrations by mediating protein-protein interactions (Ando et al., 2011). In the absence of a rigorous phylogeny, the observation of a widespread occurrence of this motif throughout the RNR family and other proteins had suggested that this domain is evolutionarily mobile (Aravind et al., 2000). However, little is known about how RNRs may have gained and lost ATP-cones during evolution. Detailed analysis of ATP-cone evolution is therefore another approach that can be used to test the validity of various evolutionary scenarios for the diversification of the RNR family.

We first examined the number of copies of the domain and the homology of each with respect to the canonical motif for every sequence in our dataset (see **Methods**). As reported previously (Aravind et al., 2000; Jonna et al., 2015), we find the motif in classes I, II, and III. In our tree, 48.6 and 48.7% of sequences in the class III and class I clades, respectively, contain ATP-cones. Both classes have subclades with multiple ATP-cones, with a particularly high occurrence rate in certain class I subclades. The class II clade has the lowest distribution of ATP-cones with 11.3% of sequences containing the domain and no sequences containing multiple copies. Although the low distribution of domains in class II has been reported before (Jonna et al., 2015), the presence of any ATP-cone domains in class II sequences remains surprising because these enzymes do not require additional subunits for radical generation and are thus not known to utilize protein-protein interactions to modulate activity. Organisms that host an ATP-cone-containing class II RNR include the anaerobic thermophile *Vulcanisaeta moutnovskia* and the aerobic ammonia-oxidizing bacterium *Nitrosococcus halophilusis*. Only one class II RNR with an ATP-cone domain has been studied (that of *Thermoplasma acidophilum*), for which activity regulation was not observed, although four dATP molecules were found to bind each α dimer (Eliasson et al., 1999). As expected, the ATP-cone motif is absent from the minimal class Ø sequences.

We next isolated ATP-cones sequences and calculated an SSN (**Figure 5A**). The major clusters were colored and mapped onto the full RNR α phylogeny (**Figure 5B**). Interestingly, we observe a large cluster of homologous ATP-cone sequences (**Figure 5A**, purple nodes) that map over the entire phylogeny, covering all three classes of RNRs (**Figure 5B**, purple color strips). Moreover, class II and III ATP-cones remain clustered together at even more stringent alignment score cutoffs. This unexpected homology suggests a possible shared ancestry and a single origin of the ATP-cone domain in the last common ancestor (LCA) to the modern RNR. The presence of ATP-cones in the class II clade can then be explained by the retention of the ATP-cone domain from the LCA. These observations support a model in which the ATP-cone was fixed in the evolutionary trajectory of the class I and III RNRs and lost in most class II RNRs as a separate subunit is not required. It is important to note that the absence of ATP-cones in the class Ø clade does not preclude the LCA of the class Ø, I, and II from having had an ATP-cone.

**Figure 5.**
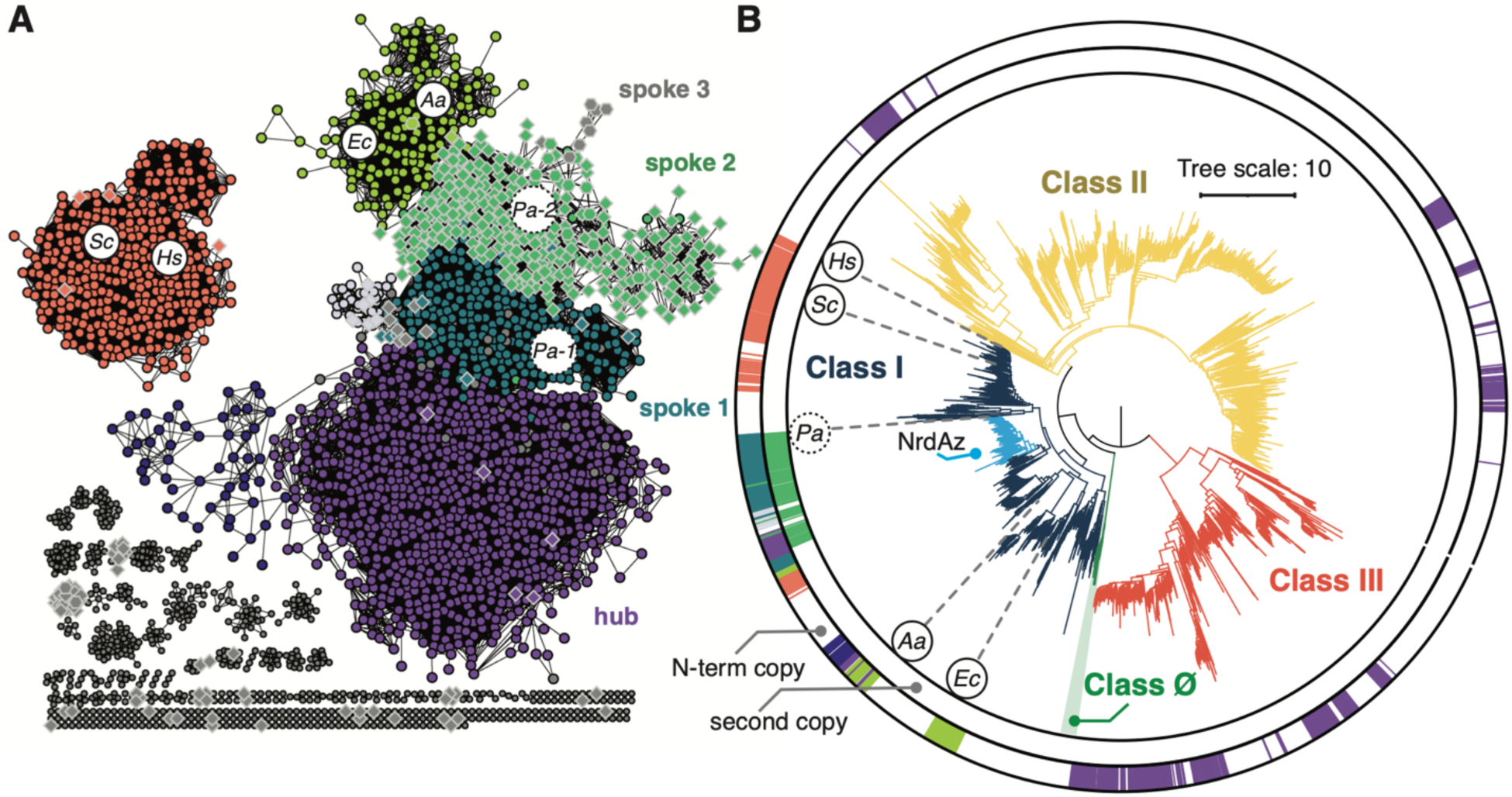
The ATP-cone domain likely originated from a single origin. (**A**) A sequence similarity network (SSN) of only ATP-cone domains extracted from 2653 ɑ sequences in our dataset. Each node is a single ATP-cone sequence, and each connecting edge indicates sequence homology of greater than E^-20^ between nodes, corresponding to a minimum identity of 34% over the entire sequence. Circular nodes correspond to the N-terminal ATP-cones, diamond nodes with gray outlines correspond to second copies in tandem ATP-cone domains. Hexagons with gray outlines correspond to the third copies in tandem. Gray-filled nodes are not represented on the circumference of the tree in panel B for clarity. (**B**) Phylogenetic tree from Figure 2 labeled with color strips corresponding to colors of the major SSN clusters in panel A. The outer color strips correspond to N-terminal ATP-cones (circular nodes in SSN), and the inner color strips correspond to the second copies in tandem ATP-cones (diamond nodes in SSN). Known ATP-cone containing sequences are labeled with organism ID (PDB accession code in parentheses): *Ec, Escherichia coli* (5cnv); *Aa, Aquifex aeolicus* (7agj); *Pa, Pseudomonas aeruginosa* (5im3); *Sc, Saccharomyces cerevisiae* (1zyz); *Hs, Homo sapiens* (3hnc). The exact sequence for the organism ID in the dashed circle is not on the tree, however its approximate location is represented by sequences with ≥ 80% sequence identity.

Notably, we observe a similar pattern of homologous clustering among individual domains within tandem ATP-cones, many of which are found in the class I NrdAz clade (**Figure 5B**). Additionally, when we examine the location of these individual copies in the SSN, we observe a “hub-spoke” topology. The most N-terminal copies within tandem ATP-cones are found in a cluster (**Figure 5A**, teal circular nodes in spoke 1) that extends from the central “hub” (**Figure 5A**, purple circular nodes) like a “spoke”, indicating that they share ancestry with the single ATP-cones found in all three classes. In the majority of cases, the second copies within these tandem ATP-cones are found in a cluster (**Figure 5A**, light green diamond nodes in spoke 2) that further extends from the cluster of N-terminal copies. Likewise, the third copies (**Figure 5A**, gray hexagonal nodes in spoke 3) extend out of the cluster of second copies. Previously, it was demonstrated with the two tandem ATP-cones of *Pseudomonas aeruginosa* class Ia RNR that only the N-terminal copy is functional, while the second copy is unable to bind ligands (Johansson et al., 2016a). Based on this finding, it was proposed that the N-terminal copy was acquired to regain allosteric regulation after the inner copy had lost function due to sequence degradation. By mapping sequence conservation onto predicted structures of tandem ATP-cones, we find that the ligand-binding residues, which are highly conserved in N-terminal ATP-cones, are indeed more variable in the second and third copies (**Figure 5 – figure supplement 1**). However, our SSN supports a converse hypothesis that a gene duplication event occurred with an active ATP-cone sequence, and the inner copies, no longer necessary for regulation, then lost function.

Two other notable class I clusters are observed. These include an island of single ATP-cones from eukaryotic class I RNRs (**Figure 5A**, salmon nodes) and a cluster of single ATP-cones from bacterial class I RNRs (**Figure 5A**, yellow-green nodes) that extends from the central hub. The clustering of *E. coli* and *Aquifex aeolicus* ATP-cones is interesting in light of recent studies (Funk et al., 2021; Rehling et al., 2022) that report structures with two ATP molecules bound in the domain in a similar fashion. The only other ɑ subunit structure with two nucleotides bound in an ATP-cone is that of the N-terminal domain of *P. aeruginosa* class Ia RNR. In this structure, one dATP is bound in a site that appears to be shared among all ATP-cones, while the second dATP is in a slightly different location than the second ATP site observed in the *E. coli* and *A. aeolicus* structures. Thus, these differences in ligand binding (or the lack thereof, in the case of nonfunctional ATP-cones) may be reflected in the clustering within the SSN.

### The class I phylogeny provides a parsimonious model for the loss and gain of ATP-cones

To examine possible evolutionary trajectories for the loss and gain of ATP-cones, we next investigated the class I RNRs, which display the greatest diversity in this motif. Subclassification of class I RNRs has been discussed extensively. Biochemical studies have thus far identified five subclasses of class I RNRs based on the radical cofactor used by the β subunit (Ia, Ib, Ic, Id, Ie) (Ruskoski and Boal, 2021). However, as we will discuss, biochemical classes do not necessarily correspond to phylogenetic clades. We will thus follow previously used gene-based nomenclature (with the prefix *nrd* for ribo**<u>n</u>**ucleotide **<u>r</u>**e**<u>d</u>**uctase) to describe phylogenetic clades (Lundin et al., 2009). In the most recently published phylogeny consisting of 342 representative class I ɑ sequences with 26 class II sequences as the outgroup, 10 clades were identified (e.g., NrdAe, NrdAg, etc.), but many were part of an unresolved polytomy (Martínez-Carranza et al., 2020).

Our RNR ɑ phylogeny includes 2023 class I ɑ sequences from 369 viruses, 1547 cellular organisms, and 24 ecological metagenomes. We resolve 11 deep-branching clades, including a newly observed viral NrdAe clade (**Figure 6A)**. The majority of nodes display high branch supports with the notable exception being that of NrdAn and NrdAy/NrdAz. The main consequence of this ambiguity is that in our tree inferences, NrdAq is the earliest diverging in group 3 (**Figure 6A)** half of the time, while in the other half, it is NrdAn. This is likely the result of the relatively small number of NrdAn and NrdAy sequences sampled (68 and 7, respectively, in our dataset). With the exception of this ambiguity, the topology shown in **Figure 6** is the most representative (8/20 inferences) (**Figure 6 – figure supplement 3**). In group 2, the modest branch support between NrdE and NrdAh results in some ambiguity on whether NrdAg or NrdAh is the most ancestral. Nonetheless, because of our large sequence dataset and the use of a structural alignment, there are no unresolved polytomies. Thus, our results show significantly improved confidence in the topology of the class I phylogeny.

**Figure 6.**
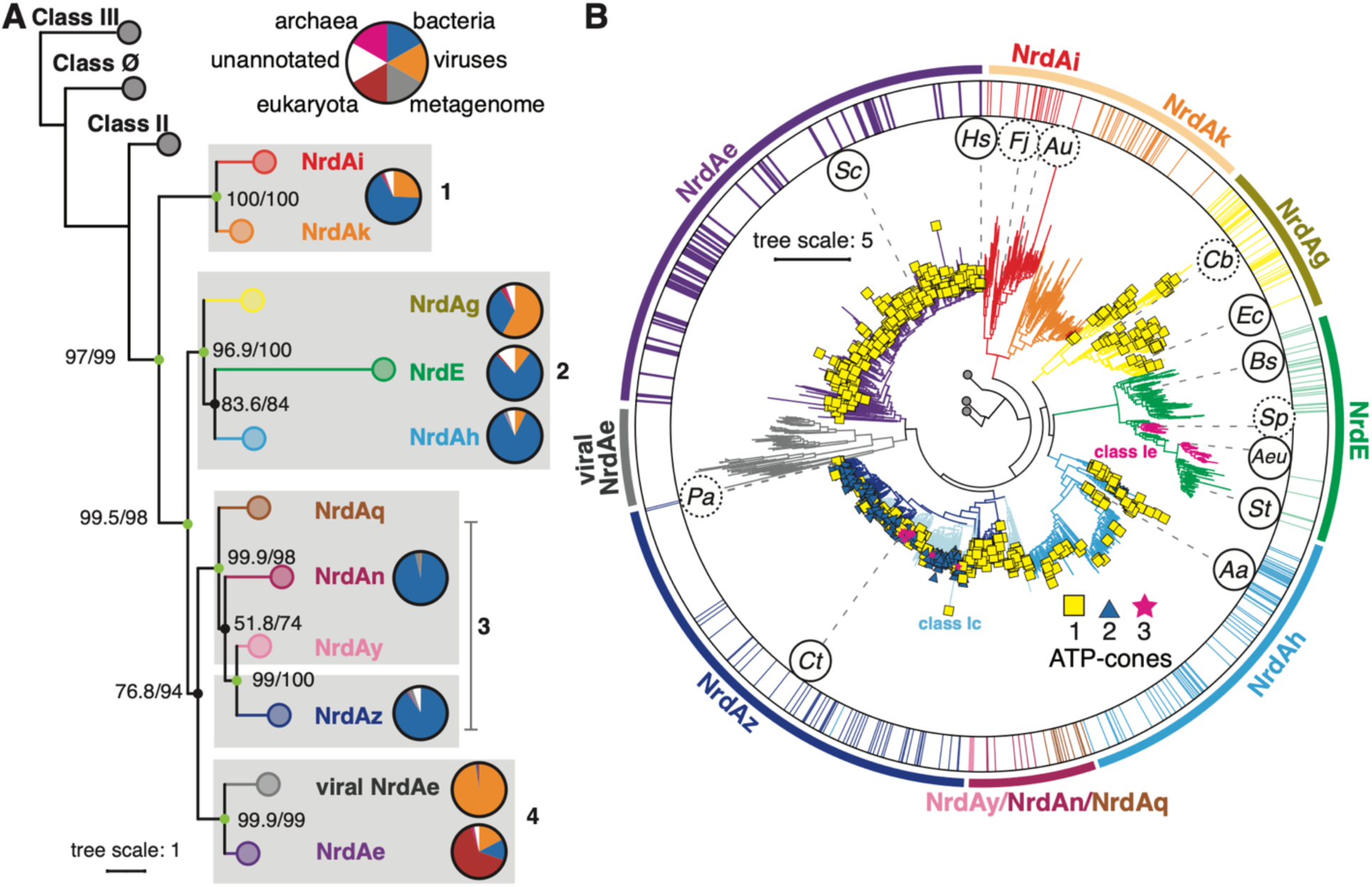
Topology of the class I RNR clade provides a parsimonious model for the loss and gain of ATP-cones. (**A**) Collapsed view of the class I clade pruned from the tree in Figure 2. The tree is labeled with SHaLRT/UFboot2 supports on important nodes. Well supported branches (SH-aLRT ≥ 80% and UFboot ≥ 95%) are shown as light green nodes, supports under this are black. Each subclade is labeled with gene based nomenclature as described in (Martínez-Carranza et al., 2020). The branches are clustered and numbered for reference in the text. The pie charts to the right of each clade show the distribution of superkingdom: viruses (orange), bacteria (blue), sequences from environmental metagenomes (grey), eukaryote (ref), archaea (magenta), unannotated in UniProt as of Jan 2022 release (white). (**B**) The expanded class I clade, colored as in panel A. In group 3 sequences (NrdAq, NrdAn, NrdAy, NrdAz), branches shown in light blue correspond to putative class Ic ɑ sequences, which are associated with a β subunit that does not have a tyrosine at the expected radical site; otherwise they are colored in dark blue. Branches in NrdE colored in magenta are putative class Ie sequences. Tips of branches are marked by the number of ATP-cones on the sequence if present: one domain (yellow squares), two domains (blue triangles), three domains (magenta stars). The organism IDs enclosed by circles are previously studied class I sequences discussed in the text. Organism IDs in dashed circles are not on the tree but their locations are approximated by sequences with ≥ 80% sequence identity. The outer color strips represent the locations of sequences used in the previous phylogeny published by Martínez-Carranza et al., 2020. Organism IDs in clockwise order: *Hs, Homo sapiens; Fj, Flavobacterium johnsoniae; Au, Actinobacillus ureae; Cb, Clostridium botulinum; Ec, Escherichia coli*; *Bs, Bacillus subtilis; Sp, Streptococcus pyogenes; Aeu, Aerococcus urinae; St, Salmonella typhimurium; Aa, Aquifex aeolicus*; *Ct, Chlamydia trachomatis; Pa, Pseudomonas aeruginosa*; *Sc, Saccharomyces cerevisiae*. Classes II, III, and Ø are shown collapsed at the root.

When compared with the most recently published class I ɑ phylogeny (Martínez-Carranza et al., 2020), the most striking difference we observe is the placement of the NrdAi/Ak clades versus the NrdE clades (**Figure 6** – figure supplement). In the previous phylogeny, the NrdE clade is the first to diverge from the LCA of class I, whereas in our analysis, we observe the NrdAi/Ak clades diverging first (**Figure 6A**, group 1).

These two, largely bacterial clades consist of the shortest sequences in the class I ⍺ phylogeny: sequences in the NrdAi clade have a median length of 564 amino acids and include biochemically characterized class Id RNRs, such as those of *Flavobacteria johnsoniae* and *Actinobacillus ureae* (Rose et al., 2019) (**Figure 6B**, *Fj* and *Au*), while the NrdAk clade consists of yet-uncharacterized sequences (**Figure 6B**) having a median length of 620 amino acids. This early divergence of NrdAi/Ak sequences is supported by the majority of our topologies (9/10 inferred with LG+R10; 8/10 inferred with WAG+R10), whereas only one topology reconstructs the class Ib RNRs as ancestral (**Figure 6 – figure supplement 3**).

In the majority of our inferences (13/20), the NrdE clade instead emerges from a largely bacterial group of sequences that includes NrdAg and NrdAh (**Figure 6A**, group 2). The NrdAg clade includes biochemically characterized class Ia sequences with a single ATP-cone (median length of 764 aa), such as the best-studied enzyme from *E. coli* (**Figure 6B**, *Ec*), whereas the NrdAh clade (median length of 780 aa) includes the *A. aeolicus* class Ia RNR (**Figure 6B**, *Aa*), which, as described earlier, contains an ATP-cone that displays homology with that of the *E. coli* enzyme. Although there is some ambiguity in whether NrdAg or NrdAh diverges first from group 2, the divergence of the NrdE clade from single ATP-cone-containing bacterial sequences is interesting as this clade is largely associated with the class Ib sequences. Class Ib ɑ sequences are notable for having only half of the ATP-cone motif at the N-terminus (Parker et al., 2018), and thus, they are shorter (median length of 718 aa) and generally lack activity regulation (Eliasson et al., 1996; Jordan et al., 1994). We observe two major subclades within the NrdE clade (**Figure 6B**), each with one structurally characterized sequence. One subclade contains the *Bacillus subtilis* class Ib RNR (**Figure 6B**, *Bs*), which was recently shown to have evolved a convergent form of activity regulation involving its partial ATP-cone (Thomas et al., 2019), while the other subclade includes the *Salmonella typhimurium* class Ib RNR (**Figure 6B**, *St*), which does not have activity regulation (Jordan et al., 1994). Interestingly, sequences homologous to the previously studied class Ie RNR appear as small groups that emerge out of this second NrdE subclade (**Figure 6B,** magenta sequences). Class Ie RNRs utilize a dihydroxyphenylalanine (DOPA) radical in the β subunit and are currently the only known metal-independent class I RNRs (Blaesi et al., 2018; Srinivas et al., 2018). The placement of the class Ie RNRs within the NrdE clade demonstrates that the mechanism of radical generation can vary within a subclade.

The other half of the class I phylogeny consists of 1127 sequences (**Figure 6A**, groups 3/4). Group 3 is comprised of largely bacterial RNRs (**Figure 6A**) and includes NrdAq/An/Ay/Az. Unlike the rest of the class I phylogeny, group 3 contains many ⍺ sequences for which the corresponding β subunit contains a redox-inert residue rather than a tyrosine as the expected radical site. Many of these class Ic-like sequences are found in the NrdAz and NrdAq clades (**Figure 6B**, light blue branches). The NrdAz clade is also notable for having many subclades and a high occurrence of sequences with multiple ATP-cones, such as the *P. aeruginosa* class Ia RNR and *Chlamydia trachomatis* class Ic RNR, which have 2 and 3 ATP-cones, respectively (**Figure 6B**, *Pa* and *Ct*) (Johansson et al., 2016b; Roshick et al., 2000; Torrents et al., 2006). As discussed in our SSN analysis of ATP cones (**Figure 5B**, NrdAz), the N-terminal copy within the tandem ATP-cones in NrdAz sequences is most homologous to the ancestral ATP-cone motif. Finally, in group 4, we observe one predominantly eukaryotic clade with the majority of sequences having a single ATP-cone (**Figure 6**, NrdAe) as a sibling clade to a previously unobserved clade of eukaryote-associated viruses that lack ATP-cones (**Figure 6**, viral NrdAe).

Combined with our SSN analysis (**Figure 5**), our class I ɑ phylogeny (**Figure 6**) provides a parsimonious model for the loss and gain of activity regulation. The trajectory we observe in our phylogeny supports the idea that the LCA of the class I RNRs had a single ATP-cone. Based on this model, the short NrdAi/Ak sequences (**Figure 6A**, group 1) were the first to diverge with the loss of the ATP-cone. A single ATP-cone was largely retained in the bacterial RNRs in group 2. Class Ib RNRs can then be explained as having emerged from this clade (**Figure 6**, NrdE) as a result of gene duplication in bacteria, in which the N-terminal ATP-cone was truncated. Interestingly, evidence for evolutionary pressure to compel activity regulation can be seen in RNRs lacking ATP-cones. One example is the *B. subtilis* class Ib RNR as described above, where new ligand-binding sites appeared to have evolved within the truncated ATP-cone motif (Thomas et al., 2019). Additionally, some class Id and Ib RNRs have been reported to contain ATP-cones on the β subunit, which could be a result of a different strategy that became fixed from similar evolutionary pressure (Rozman Grinberg et al., 2018). In our dataset, we find that ATP-cones exist on the β subunit almost exclusively where the operon is organized such that α is transcribed downstream from β (**Figure 6 – figure supplement 2**), contrasting with the majority of the class I RNR operons, which are organized with α upstream from β. We thus hypothesize that the β subunit acquired an N-terminal ATP-cone through gene insertion in the middle of ATP-cone-containing α sequence. We observe an especially high occurrence of operons organized as β-α in the NrdAi (class Id) clade (70/104 sequences). Interestingly, the latest diverging NrdAi subclade is predominantly associated with ATP-cones on the β subunit. In group 3, we observe a high occurrence of multiple ATP-cones in the NrdAz clade (**Figure 6B**, blue triangles or magenta stars at branch tips), which we hypothesize to have initially been produced by gene duplication of a functional ATP-cone based on our SSN analysis.

Finally, group 5 is noteworthy in that it provides insight into host-virus relationships and their possible influence on the loss and gain of ATP-cones. While eukaryotic RNRs (NrdAe) have retained a single ATP-cone, viral RNRs associated with eukaryotes (viral NrdAe) appear to have lost the ATP-cone. The sequence similarity of the NrdAe and viral NrdAe RNRs is likely derived from gene exchange between viruses and hosts (Weiss, 2017), while the loss of the ATP-cone could be the result of evolutionary pressure to maintain a compact genome (Belshaw et al., 2007). This observation can also explain the loss of ATP-cones in the class Ø clade, which are associated with cyanophages.

### Long class II C-termini contain folded domains

One of the most unusual features of the class II RNRs is the presence of monomeric enzymes that structurally mimic the allosteric dimer interface (Sintchak et al., 2002). Based on analysis of the insertion length between βB and ɑB (**Figure 1A**, loop 1) and structure prediction by AlphaFold2, we first identified the locations of monomeric and dimeric class II RNRs in our phylogeny. Sequences predicted to be monomeric (29-82 residue insertion) were found to cluster largely within a single clade (**Figure 7A**, orange color strips). The remaining majority (2967 out of 3347 sequences in our dataset) were predicted to be dimeric. To further corroborate these assignments, we examined another unique feature of class II RNRs. Unlike class I and III RNRs which are specific for ribonucleoside diphosphates (NDPs) and triphosphates (NTPs), respectively, class II RNRs show diversity in substrate preference. Substrate preference is associated with a 4-5 residue sequence motif preceding the catalytic barrel (**Figure 1A**, left star), with PNSP (NDP-specific) and PAGR (NTP-specific) being the two most predominant patterns (Schell et al., 2021; Wei, 2015). The two class II RNRs for which structures have been solved represent each of these cases: the monomeric class II RNR of *Lactobacillus leichmannii* is specific for NTPs (Sintchak et al., 2002), while the dimeric class II RNR of *Thermotoga maritima* is specific for NDPs (Larsson et al., 2010). In 17 out of our 20 inferences, the monomeric clade is placed near the root of class II RNRs along with other NTP-specific clades, while the later diverging clades are dominated by sequences that are predicted to be NDP-specific (**Figure 7 – figure supplement 1**). This observation suggests that the monomeric class II RNRs diverged early on from a dimeric ancestor. Such a scenario is consistent with our evolutionary model that the class I and II RNRs diverged from an ancestor to the class Ø RNRs, which, based on our structural results, are likely to exist as dimers in the presence of intracellular nucleotides.

**Figure 7.**
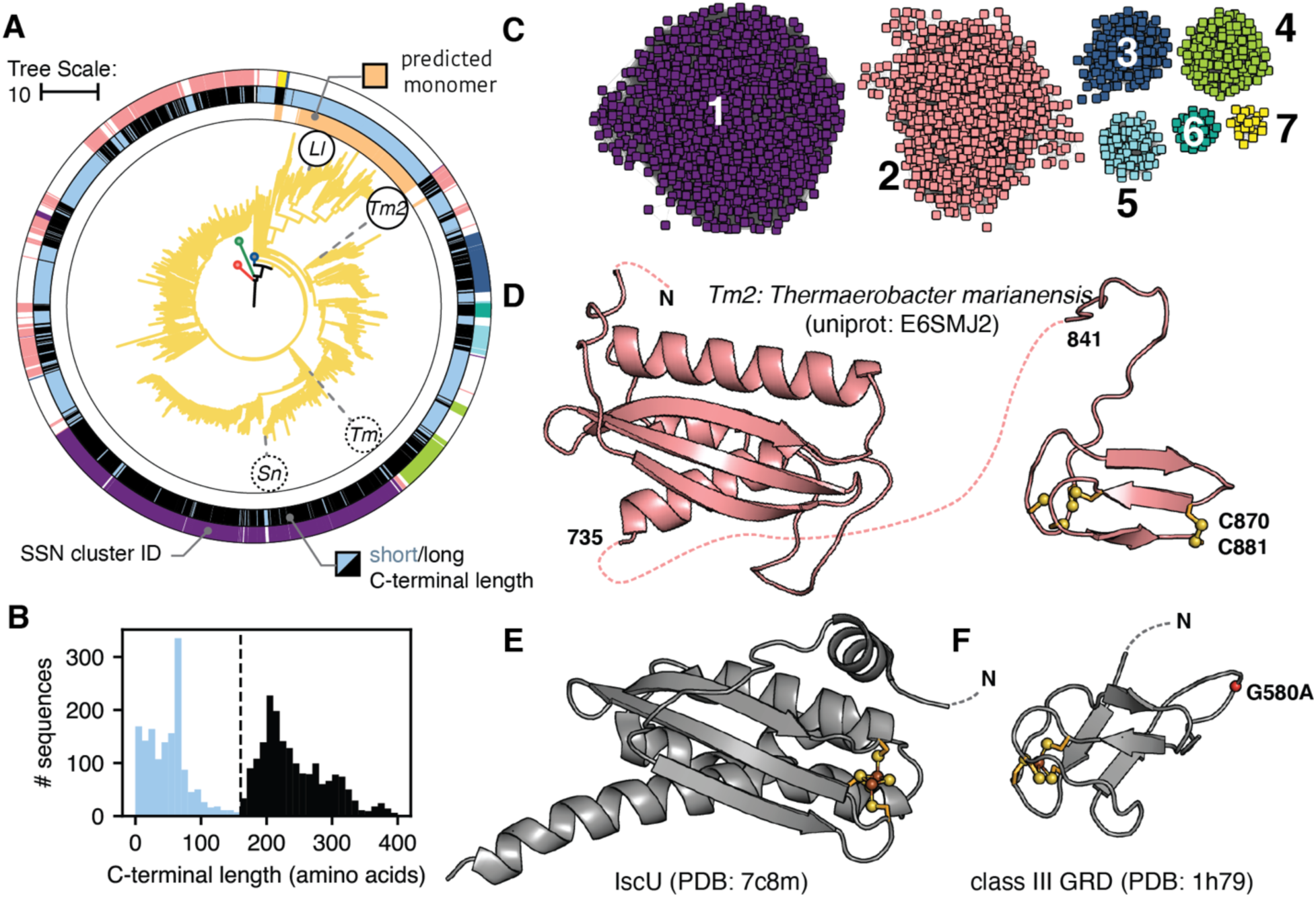
Diversity of the class II sequences is found in the C-terminal tail. (**A**) Class II clade pruned from tree in Figure 2. Sequences predicted to represent monomeric class II RNRs are shown as peach color strips in the innermost ring of the circumference. The middle color strips correspond to the length of the C-terminal tails, where light blue is short and black is long, as depicted in panel B. The outer color strips correspond to the SSN clusters shown in panel C. (**B**) The C-terminal length distribution is bimodal. (**C**) An SSN of the C-terminal sequences. Only the largest seven clusters with the most sequences are shown for clarity. (**D**) Structure prediction of a representative C-terminal sequence (*Thermoaerobacter marianensis*) from cluster 2 (pink) in panel C. The left domain shows structural homology to the IscU protein. The right domain is a zinc finger. (**E**) Crystal structure of IscU protein (PDB: 7c8m). (**F**) Glycyl radical domain of the class III RNR (PDB: 1h79). In panels D-F sulfur and iron are colored in yellow and brown, respectively. In panel F, the glycine residue associated with radical formation was mutated to an alanine and the Cα carbon is shown as a red sphere.

We next examined additional characteristics of the class II RNRs and discovered surprising diversity in the C-terminus after βJ, the last common fold of the catalytic barrel. Unlike the C-termini of class I and III RNRs, class II C-termini span a wide range of lengths, and the distribution is unexpectedly bimodal (**Figure 7B**). The first peak in the distribution corresponds to sequences with short C-termini (≲ 160 residues, a length similar to those of class I and III RNRs), while the second peak corresponds to sequences with unusually long C-termini (160-400 residues). The majority of the monomeric class II RNRs and many of the dimeric enzymes have short C-termini (**Figure 7A**, middle color strips in light blue). Sequences with long C-termini are exclusively dimeric class II RNRs except for two minor subclades in the monomeric clade (**Figure 7A**, middle color strips in dark blue).

In the case of the best-studied class II RNR, the monomeric enzyme from *L. leichmanii*, it was shown that the double cysteines on its short C-terminus serve to reduce the active site after each turnover, much like in class I RNRs (Booker et al., 1994). However, the function of long C-termini, which are present in a large subset of class II RNRs (56% in our tree), is not well understood. Studies of class II RNRs from *T. maritima* (Eliasson et al., 1999; Larsson et al., 2004) and *Stackebrandtia nassauensis* (Loderer et al., 2017) have suggested that long C-termini may be involved in oligomerization. Through a combination of homology modeling and metal-binding assays, it was further proposed that the six cysteines at the very C-terminus form a zinc finger domain that facilitates the reduction of the active site, in a manner reminiscent of the C-terminal double cysteines in class I RNRs. Although the endogenous reductant remains unknown, activesite reduction by the C-terminus was supported by a comparison of small-molecule reductants (Loderer et al., 2017).

To explore the classification and evolution of class II C-termini, an SSN was constructed with the isolated C-terminal sequences (**Figure 7C**), and the seven largest clusters were mapped onto our tree (**Figure 7A**, outer color strips). Using AlphaFold2 (Jumper et al., 2021), we predicted the structure of a representative sequence from each of the largest SSN clusters, all of which consisted of long C-termini. Two recurring structural motifs are observed (**Figure 7D**). Surprisingly, the first motif is a domain with an α+β fold (**Figure 7D**, left) that shows structural homology with the iron-sulfur cluster assembly protein IscU when searched against the PDB (**Figure 7E**). The second motif at the very C-terminus is a zinc finger domain (**Figure 7D**, right) as predicted previously by homology modeling (Loderer et al., 2017). The zinc finger domain is found in the majority of dimeric class II RNRs (71%) with either a long or short C-terminus but is only found in the two minor subclades of monomeric class II RNRs with long C-termini.

A role for the putative IscU-like domain has not been reported in the RNR literature. However, in the iron-sulfur cluster assembly pathway, IscU acts as a scaffold for the maturation of a [2Fe-2S] cluster within the monomer or a [4Fe-4S] cluster through the dimerization of two monomers (Ayala-Castro et al., 2008). The coordination of Fe-S clusters in IscU involves two cysteines and an aspartate, with the additional ligand being either a cysteine or histidine depending on the species. Although the majority of IscU-like motifs in class II RNRs do not contain any cysteines at the equivalent positions, 253 out of the 600 sequences in cluster 2 of the SSN (**Figure 7A and C,** salmon color strips and nodes) contain cysteines at these four positions, indicating possible ancestry of these residues from an IscU-like protein. Furthermore, cluster 2 sequences are found in distantly related subclades (**Figure 7A**, salmon color strips), suggesting that this C-terminal motif may have shared a common origin. We hypothesize that this domain was acquired through insertion of an IscU-like protein, which later evolved to support other functional needs of class II RNRs. The predicted structures of class II IscU-like domains are relatively conserved, with insertion or duplication occurring in SSN clusters 5, 6, and 7 (**Figure 7 – figure supplement 2**). Although class II RNRs are not known to utilize Fe-S clusters, dimerization of the IscU-like domain may explain the higher order oligomerization previously reported for the dimeric class II RNRs from *T. maritima* and *S. nassauensis* (Eliasson et al., 1999; Loderer et al., 2017).

All predictions of the C-terminal zinc finger domain are structurally identical, with four cysteines poised to coordinate a Zn^2+^ ion and a cysteine pair located at the opposite end (**Figure 7D**, right). This domain is reminiscent of the zinc finger in the C-terminal glycyl radical domain of class III RNRs (**Figure 7F**). In class III RNRs, the zinc finger is thought to provide structural stability for the glycyl radical (Logan et al., 2003), which is located on a C-terminal loop. Although the function of the zinc finger domain in class II RNRs remains to be explored, it may play a similar structural role by positioning the cysteine pair for activesite reduction.

C-terminal sequences without a zinc finger motif were also examined. The majority of these (64%) contain two cysteines similar to class I RNRs, with smaller populations containing more than two cysteines (6%) or only one cysteine (4%). The number of residues between the double-cysteine motif is, however, highly variable, ranging from no residue to more than 4 residues. In contrast, most class I RNRs show a consensus CXXC motif at the C-terminus, with a minor population having a CXXXXC motif such as the well-studied *E. coli* class Ia RNR. In addition, 26% of C-terminal sequences without a zinc finger domain contain no cysteines. Roughly 40% of these represent a subclade of split class II RNRs, which consist of two chains, NrdJa and NrdJb. An AlphaFold prediction of a representative species shows that NrdJb contains the IscU-like and zinc finger domains described above for other long C-termini, while NrdJa contains the catalytic barrel and an extra folded domain at the very C-terminus, which may be involved in facilitating interactions with NrdJb. The remaining sequences with no C-terminal cysteines may possibly represent enzymes that utilize small-molecule reductants that can directly reduce the active site, although biochemical studies are needed to corroborate such behavior.

### Class III RNRs can be classified by the finger-loop motif

Class III RNRs are the most distantly related to the other classes both in terms of sequence homology and the oxygen sensitivity of the radical cofactor (**Figure 2-3**). Structures of two class III RNRs have been reported thus far, those from T4 phage (Andersson et al., 2000; Logan et al., 1999) and *T. maritima* (Aurelius et al., 2015; Wei et al., 2014a). These structures reveal key features that distinguish class III RNRs from class Ø, I, and II RNRs, the most apparent being the unique dimer orientation facilitated by a long helical insertion in place of loop 1 (**Figure 1D**). The structures also capture the C-terminal glycyl radical domain (GRD) positioned within the active site, such that the glycyl radical site is poised for radical transfer with the residue at the tip of the finger loop. In structures of the T4 phage class III RNR, the finger loop contains a single cysteine at the tip (**Figure 8 – figure supplement 1A**). This placement is consistent with all existing structures of class I and II RNRs and supports our current understanding that nucleotide reduction is initiated by a thiyl radical. However, in structures of the *T. maritima* class III RNR (solved independently by two research groups) (Aurelius et al., 2015; Wei et al., 2014a), an isoleucine is instead found at the tip of the finger loop, placing the expected cysteine at the back of the catalytic barrel in an unmodeled region that contains a double cysteine motif. Making sense of this finger-loop conformation has been an outstanding problem involving the class III RNRs.

**Figure 8.**
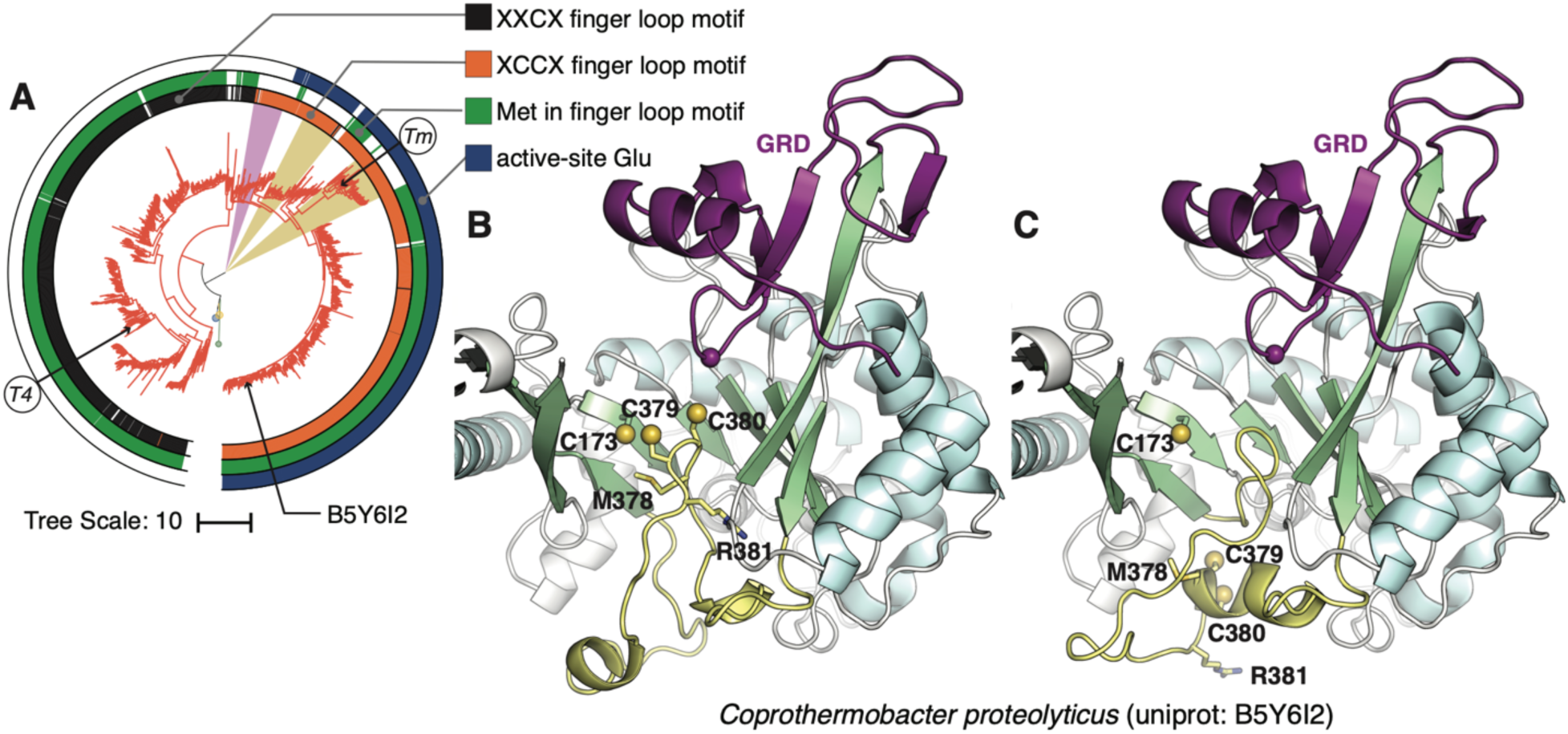
Class III diversity is driven by active-site and finger loop motifs. (**A**) Class III clade pruned from tree in Figure 2. The innermost colors trip defines the two major subclassification of class III α subunits based on the number of cystines in the finger-loop motif: single-cysteine (black) and double-cysteine (orange). The middle color strips are green if the finger-loop motif begins with a methionine. The outer color strips are blue if the active site contains a glutamate as a putative proton source. The gold wedges highlight clades where a glutamate is present but not a methionine in the finger-loop motif, characteristics associated with only using thioredoxin as the reductant. The purple wedge highlights the subclade that neither contains a glutamate nor a methionine; these sequences have been previously subclassified as NrdD3 and thought to employ a novel reduction mechanism. Sequences with known structures are labeled as *T4, Enterobacteria* phage T4 (PDB 1h7a) and *Tm, Thermotoga maritima* (PDB 4ue3). (**B-C**) Slice-through views of predicted models of the class III α from *Comprothermobacter proteolyticus* (UniProtID B5Y6I is mapped in panel A). The finger-loop is colored yellow, and the cysteine sulfur atoms are shown as spheres. The glycyl radical domain (GRD) is shown in purple. In panel B, the finger-loop is predicted to have both cysteines on the tip, whereas in panel C, the finger loop is in the same conformation as in the *T. maritima* structure with the cysteines at the back of the barrel.

Based on small-scale class III RNR bioinformatic analyses, it was shown that the finger loop region of class III RNRs contains distinct sequence motifs (Funk, 2015; Wei et al., 2014a). In our dataset of 1375 class III ⍺ sequences, we find there are two dominant motifs: a single-cysteine motif (MGCR) or a double-cysteine motif (MCCR), where the methionine is 97% and 83% conserved in the respective motifs. The catalytic cysteine is the third residue in these motifs. When mapped onto our phylogeny, sequences with double cysteines cluster tightly, and their divergence from those with the single-cysteine motif is monophyletic, where splitting occurs at a single node (**Figure 8A**, black vs orange color strips). This suggests that the two currently existing structures may be representative of two subclasses of class III RNRs.

To explore possible conformations of the finger loop, we employed AlphaFold2 (Jumper et al., 2021) to predict structures of class III RNRs with a double-cysteine motif. For the *T. maritima* class III sequence, AlphaFold2 predicts structures with the missing region in the finger loop modeled (**Figure 8 – figure supplement 1B**), but otherwise identical to the crystal structure. This is expected, as the machine-learning model enforces agreement with existing structures in the training dataset. Notably, however, the confidence score for the finger-loop prediction is low (**Figure 8 – figure supplement 1C**). When used to predict structures of more distant class III RNRs with double cysteines (i.e., not within the same subclade as the *T. maritima* enzyme), several or all five models generated by AlphaFold2 (per sequence) predict the double cysteines at the tip of the finger loop (**Figure 8B)**, while the remaining models predict the same conformation as the *T. maritima* crystal structure (**Figure 8C)**. The finger-loop region, when defined as the insertion between βE and βF (**Figure 1A**), is relatively long in all RNRs (∼40 residues in class III RNRs) and often contains secondary structure, typically a small helix preceding the catalytic cysteine (**Figure 1A**, yellow). Comparison of the two finger-loop conformations observed in AlphaFold predictions suggest that changes in secondary structure may facilitate the movement of the double cysteines to the back of the catalytic barrel (**Figure 8B-C**).

Biochemical studies have identified two mechanisms of active-site reduction in class III RNRs that appear to correlate with the number of cysteines at the tip of the finger loop (Wei et al., 2014b, 2014a). In class I and II RNRs, a conserved cysteine pair on the βA and βF strands provide two electrons during the reductive half-reactions. However, in class III RNRs, only the cysteine in βA is highly conserved (92.6%). It was previously shown that in *E. coli* class III RNR, the methionine in its single-cysteine MGCR motif forms a thiosulfuranyl radical with the conserved cysteine on βA to donate one electron, which subsequently is reduced by formate (Wei et al., 2014b). Such a catalytic role would explain the fact that the methionine is conserved (97%) in this subclass of enzymes. In the case of *Neisseria bacilliformis* class III RNR, it was shown that one of the consecutive cysteines in its SCCR motif (lacking the upstream methionine) forms a disulfide bond with the βA cysteine in a manner similar to class I and II RNRs to donate two electrons, which can then be reduced by thioredoxin (Wei et al., 2014a). However, from our analysis, the majority (83%) of class III sequences with double-cysteine motifs contain a methionine at the start of the motif and thus appear to contain essential components for utilizing either formate or thioredoxin as reducing equivalents. When mapped onto our tree, the methionine-containing sequences appear to be ancestral, consistent with the usage of formate as an electron donor in anaerobic metabolism (**Figure 8A**, green color strips). As noted previously (Mulliez et al., 1995), the use of formate links the class III RNRs with the glycyl radical enzyme (GRE) pyruvate-formate lyase and suggests a common origin for the RNR and GRE families.

In the study of *N. bacilliformis* class III RNR (Wei et al., 2014a), it was further proposed that an active-site glutamate that is at an equivalent position to the established proton source in *E. coli* class Ia RNR (in the reduction of the 3′-keto-deoxynucleotide intermediate) is essential for formate-independent nucleotide reduction. By comparing where sequences with this glutamate and the finger-loop methionine map on our tree, we identified two small subclades, one of which includes the enzymes from *N. bacilliformis* and *T. maritima*, that contain glutamate but do not have methionine and are thus likely to solely utilize thioredoxin as the reductant (**Figure 8A**, orange wedges). We also observed the clade previously classified as NrdD3 that contains a QCCR/A motif with neither the methionine nor glutamate (**Figure 8A**, purple wedge), which has been suggested to employ a novel reduction mechanism (Wei et al., 2014a). We note that additional cysteines exist on the β strands near the finger loop in many class III RNRs. For instance, 33% of the class III sequences contain two consecutive cysteines on βA, which is not correlated with the number of cysteines on the finger loop. These additional cysteines may have significance in the reductive half-reaction of RNRs.

## Discussion

In this study, we were able to perform a large-scale phylogenetic reconstruction of the highly diverse RNR family by using a structural alignment. Key bifurcations featured high branch supports and reproducibility in replicate tree inferences, allowing us to make robust interpretations. Overall, our unified RNR phylogeny showed the evolution of three major, monophyletic clades corresponding to the three biochemically known classes, with the O2-sensitive class III RNRs being the earliest to diverge. Unexpectedly, we observed a new clade, which we denoted the class Ø clade, that shares an ancestor with the LCA of the O2-dependent class I and O2-tolerant class II RNRs. This evolutionary trajectory was independently supported by evo-velocity analysis. In addition, using a combination of bioinformatic analyses and structure prediction, we observed functional diversification of the RNR family via extensions and insertions about the conserved catalytic barrel. These analyses revealed evolutionary patterns in the N-terminal ATP-cones, folded domains in class II C-termini with as-yet unknown function, and two clear lineages in the class III RNRs based on finger-loop motifs. This work highlights the expansive complexity of the RNR family and provides a roadmap for the many important questions that remain.

Evo-velocity analysis identified multiple roots in the RNR family due to the lack of intermediate phylogenetic information connecting the highly diverged class III RNRs and the other classes. Surprisingly, however, this connection was found in the N-terminal ATP-cone motifs. Previously, it was thought that specificity regulation was the only form of allosteric regulation that is common to all RNRs, while activity regulation was only present in RNRs that acquired the ATP-cone by horizontal domain transfer. Furthermore, it was suggested that the class I, II, and III RNRs independently acquired the ATP-cone domain at various points in their evolution (Aravind et al., 2000; Martínez-Carranza et al., 2020). In contrast to these previous hypotheses, our work suggests that an ATP-cone-containing ancestor to all three classes resulted in the vertical inheritance of the domain. Of the three classes, the class I ATP-cones have diversified the most, while the homology observed between the class III and II ATP-cones suggests that they may be the most similar to the ancestral form of the ATP-cone domain. Further structural characterizations of ATP-cone- containing RNRs therefore remains of great interest.

Most notably, our phylogeny revealed the novel class Ø clade. Bioinformatically, we showed that the class Ø ɑ subunit shares significant sequence homology with class II sequences, while also containing in its gene neighborhood a gene for a ferritin-like protein that resembles the class I β subunit. Structurally, we further showed that the class Ø ɑ subunit defines the minimum set of features shared by class I and II RNRs, including the dimer interface, specificity site at loop 1, and the β-hairpin. It was previously hypothesized that the LCA of RNRs may have possessed class II-like biochemistry based on the Precambrian bioavailabilities of iron and cobalt and the abiotic synthesis of porphyrin rings (Lundin et al., 2015). An alternative hypothesis is supported by our phylogeny, which places the class I mechanism as ancestral to, or having developed in parallel with, that of class II. Although we cannot classify the mechanism of radical generation in the class Ø RNRs in the absence of biochemical data, our phylogeny suggests that a class II-like enzyme resembling the extant class Ø sequences recruited a ferritin-like protein into the catalytic machinery, before diverging into class I RNRs (which retained a ferritin subunit for catalysis) and class II RNRs (which specialized in cobalamin usage). The usage of a ferritin-like protein in class Ø RNRs is supported by the observation of a stacked YY dyad in the active site of our cryo-EM structure, which suggests that long-range radical transfer is an important component of catalysis in these enzymes. Future biochemical characterization will require identification of the correct metallocofactors for activity and possible reducing partners.

Interestingly, the class Ø clade is associated with marine organisms, such as cyanophages and photosynthetic bacteria. As in the class I phylogeny, where we observed NrdAe and viral NrdAe clades evolving from a common ancestor, the presence of both viral and bacterial sequences in the class Ø clade is consistent with the dynamic exchange of genetic material between viruses and their hosts contributing to their evolution (Sullivan et al., 2006; Weiss, 2017). Additionally, viral genomes are known to be compressed in order to reduce the space they physically occupy (Belshaw et al., 2007). This genome reduction mechanism may also explain the absence of the ATP-cone in the viral NrdAe and class Ø clades. Furthermore, the lack of RNR activity regulation may reflect its nutrient-poor environment and also contribute to the fitness of the phage. Photosynthetic bacteria and phages in the ocean are particularly limited in phosphorus, which is essential for nucleotide biosynthesis (Gao et al., 2016). Studies of the *Prochlorococcus* P-SSP7 phage have shown that infection induces transcription of the phage RNR, while shutting down transcription of most host genes (Lindell et al., 2007; Sullivan et al., 2005). A notable exception was a host gene involved in RNA degradation that was upregulated by phage infection (Lindell et al., 2007). These observations, combined with estimates that sufficient deoxyribonucleotide pools for phage replication cannot be obtained by digestion of host DNA alone (Thompson et al., 2011), illustrate how nutrient-poor the environment is for these cyanophages that encode class Ø RNRs. In such an environment, allosteric inhibition of RNR activity via an ATP-cone is not only unnecessary for phages but may also be a disadvantage in their ‘arms-race’ with their hosts.

Based on our results, we present the following model for the evolution of the RNR family. The LCA of the RNR catalytic subunit likely contained an ATP-cone and shared the most biochemical resemblance to extant class III RNRs. Increasing oxygen availability selected for the divergence of the RNR family into the anaerobic LCA of class III and the oxygen-tolerant LCA of classes Ø – II, from which the minimal class Ø RNRs diverged with the loss of the ATP-cone. Class I and II RNRs sharing an ancestor would suggest that multiple strategies to adapt to the presence of oxygen were developed in parallel, rather than sequentially. Interestingly, we find sequences from the earliest diverging cyanobacterium, *Gloeobacter*, near the root of the class II clade in our phylogeny. *Gloeobacter* are notable for lacking thylakoid membranes and have been called “the missing link” between anoxic photosynthesis, which involves Fe-S chemistry, and oxygenic photosynthesis (Nakamura et al., 2003; Saw et al., 2013). Thus, along with the emergence of cyanobacteria (Schirrmeister et al., 2013), it is possible that class II RNRs evolved well before the Great Oxygenation Event (GOE), and the incorporation of an IscU-like domain in the C-terminus may be a vestige from this period. If so, this would imply that class I RNRs also emerged before the Earth’s atmosphere became permanently oxidizing. In fact, recent work suggests that oxygen availability was sufficient for the birth of O2-producing and O2-utilizing enzymes to occur well before the geochemically defined GOE (Jabłońska and Tawfik, 2021). Such an evolutionary model could explain how O2-dependent class I RNRs can share an ancestor with class II RNRs. In this respect, it is especially intriguing that class Ø RNRs are associated with cyanophages and photosynthetic bacteria.

### Outlook

In conclusion, our phylogeny presents a roadmap for future study of the RNR family. Biochemical studies of the class Ø clade will provide insight into how an ancestral radical-generating mechanism may have diverged into class I and II mechanisms. Additionally, structural and biochemical characterization of ATP-cone-containing class III RNRs may provide an understanding of shared ancestry in activity regulation. Finally, the predicted folds of the class II C-termini provide hypotheses for experimental investigation. In all, our unifying RNR phylogeny will guide future studies to fill important gaps in our ever-evolving understanding of this family of complex enzymes.

## Methods

### Sequence collection and SSN analysis

A dataset of 105,904 sequences belonging to families PF02867, PF08343, PF13597, PF00317, PF17975, and PF08471 was downloaded from the PFAM server (El-Gebali et al., 2019). An arbitrary sequence redundancy threshold of 85% was imposed on the dataset, and redundant sequences were clustered and represented as a single sequence in CD-HIT (Fu et al., 2012). Sequences that were smaller or larger than 400 or 1100 residues, respectively, were deleted from the dataset. An all-vs-all pBLAST search was performed for each sequence in the dataset using default search parameters and an E-value threshold of 10E^-240^. The search result was visualized as a force-directed graph network in Cytoscape (Hie et al., 2021; Shannon et al., 2003). The final non-redundant dataset consisted of 12,968 sequences.

### Sequence alignment

As RNR sequences have been diverging for billions of years, pair-wise sequence identity is frequently low (<15%), and homology is often undetectable between distant members of the superfamily. Conventional alignment algorithms failed to align these large and diverse datasets, which were evaluated against structural alignments of phylogenetically diverse RNRs. Instead, we employed our recently reported workflow (Spence et al., 2021) that uses ensembles of HMMs that are iteratively trained on increasingly diverse sequences until the full diversity of the dataset is captured. The initial HMM in this workflow captures residues that are tolerated at each site of the consensus RNR fold. This HMM was built from a non-redundant structural alignment of 9 diverse RNR crystal structures (1h78, 4coj, 3o0m, 6cgm, 1pem, 5im3, 3hne, 3s87, and 2x0x) that had been aligned in MUSTANG (Konagurthu et al., 2010) HMMs were built using the HMMBuild application of the HMMer suite (Finn et al., 2011). The most representative sequence from each sequence cluster in the SSN was aligned to the initial structural HMM (755 sequences). This “representative” alignment, and a corresponding maximum-likelihood (ML) phylogenetic tree that was reconstructed in IQ-TREE (Minh et al., 2020) was used as a seed alignment and guide tree, respectively, for the full alignment of all non-redundant RNR sequences. The final sequence alignment was performed in UPP from the SEPP alignment package (Nguyen et al., 2015). As this alignment workflow exploits structural information, residues that are not crystallographically resolved, such as non-conserved insertions and N-/C- terminal extensions (including the ATP cone and functional C-terminal extensions discussed in main text), are not included in the alignment. The full alignment was manually refined by removing non-conserved insertions (typically single residue insertions conserved in fewer than 1% of sequences) and poorly aligned sequences that are unlikely to be functional RNRs. The final alignment features 6,779 RNR sequences.

### Phylogenetic reconstruction

The RNR superfamily phylogeny was reconstructed by ML from the full RNR alignment. Phylogenetic inference was performed using IQ-TREE2.0 (Minh et al., 2020) on the Australian National Computing Infrastructure (NCI) GADI supercomputer and BioHPC servers at the Cornell Institute of Biotechnology. Stochastic tree-search was conducted with a stochastic perturbation strength of 0.2 to a maximum of 2000 iterations. Convergence was defined as 200 iterations of tree-search that failed to improve the current best tree. Branch supports were measured by ultrafast bootstrap approximation 2.0 (Hoang et al., 2018) and the approximate likelihood ratio test, each conducted to 10,000 replicates. The sequence evolution model, LG replacement matrix (Le and Gascuel, 2008) with a 10 category free-rate rate heterogeneity model (Kalyaanamoorthy et al., 2017), was determined by ML using ModelFinder (Kalyaanamoorthy et al., 2017) as implemented in IQ-TREE2.0 on the representative sequence alignment. To test the robustness of our hypotheses against model uncertainty, we performed an additional 10 replicates of ML tree-search using an alternate general amino-acid replacement matrix (WAG+R10) (Whelan and Goldman, 2001). The AU-test conducted to 10,000 replicates was used to compare all tree topologies (Shimodaira, 2002).

### ESM-1b evo-velocity modeling

Protein natural language processing (NLP) models can provide valuable and phylogeny-independent evolutionary insight (Alley et al., 2019; Hie et al., 2021; Rives et al., 2021). The 6,779 sequences in our dataset were embedded as high-dimensional vectors in the ESM-1b protein language model (Rives et al., 2021) and projected onto a two-dimensional directed network graph with uniform manifold approximation and projection (UMAP) (McInnes, L., Healy, J., Saul, N., & Großberger, L., 2018). Likelihoods along each edge in the network graph were computed and visualized as a vector field. Under the assumption that likelihoods inferred from the NLP model are correlated with evolutionary fitness, the vector field “flows” in the direction of evolution, and pseudotime velocity is correlated with phylogenetic depth (Hie et al., 2021).

### Analysis of ATP-cones

The N-terminal extensions align poorly with a structure-based alignment that is anchored on the core catalytic barrel. The regions N-terminal to the first common fold of the barrel were therefore aligned separately with MUSCLE and concatenated with the core-aligned sequences prior to phylogenetic inference (Edgar, 2004). The presence of ATP-cone motifs was determined by alignment against Pfam 03477 with HMMER using an E-value cutoff of 10^-3^. The domains were then mapped onto the RNR ɑ subunit phylogeny to investigate the ubiquity and evolution of the ATP-cone across phyla (Eddy, 2011; El-Gebali et al., 2019). For the SSN of the ATP cones, ATP cone sequences identified by InterPro (Blum et al., 2021) in the tree was extracted and used to construct an SSN using EFI-EST (Zallot et al., 2019). The resulting SSN was filtered with an alignment score cutoff of 20.468.

We note that a number of long class I sequences with multiple ATP-cones were misannotated as class II sequences in the UniProtKB database (Bateman et al., 2021). Upon closer examination of the *nrd* operons, these misannotated *nrdA* genes are associated with a gene for a class I radical-generating subunit (*nrdB*) immediately downstream. In our phylogeny, these sequences appear in the class I clade.

### Analysis of class II C-termini

To analyze the C-terminal extensions, the sequences from the class II clade in the tree were aligned with MAFFT (Katoh et al., 2017). A pair-wise alignment between the structurally characterized sequence of *T. maritima* class II RNR (O33839) and its closest homolog in the dataset (93% identical), *Thermotoga neapolitana* class II RNR sequence (B9K714), was performed with Clustal Omega (Sievers et al., 2011). Alignment columns after the column that corresponds to residue 639 in B9K714 (630 in O33839) were assigned as the C-termini of the corresponding sequences. The extracted C-termini sequences were used to construct an SSN using EFI-EST (Zallot et al., 2019), and an alignment score of 30 was used to filter the edges. The DALI server (Holm, 2020) was used to identify the homology of folded domains in AlphaFold2 predictions of the C-termini.

### Analysis of class III finger loop motifs

The sequences from the class III clade in the tree were aligned with MAFFT (Katoh et al., 2017). Alignment columns corresponding to residues 319 – 367 in *T. maritima* class III RNR (Q9WYL6) were assigned as the finger-loop region and extracted. Insertions in a small number of extracted sequences (22 out of 1375) were manually removed, and the sequences were realigned with MAFFT. Columns corresponding to the finger-loop motif were determined by alignment with residues 328 – 331 (SCCR) in Q9WYL6. The column corresponding to the proton donor glutamate was determined by alignment with residue 495 in Q9WYL6.

### Expression and purification of *Synechococcus* phage S-CBP4 α subunit

The gene for the α subunit of *Synechococcus* phage S-CBP4 RNR was synthesized by Genscript with a His6-SUMO (smt3) tag at the N-terminus and cloned into a pET-28c+ vector in between the *NcoI* and *XhoI* cloning sites. His6-smt3-tagged-α was grown overnight at 37 °C with 200 rpm shaking in 100 mL of LB medium supplemented with 50 µg/mL kanamycin. Large-scale growth was initiated by adding 10 mL of saturated starter culture to six 1 L volumes of kanamycin-supplemented LB medium incubating at 37 °C with 200 rpm shaking. At an OD600 of 0.4, IPTG was added to a final concentration of 0.4 mM, and cultures were shaken at 37 °C 200 rpm for 6 hours. Cells were harvested by centrifugation at 4 °C (15 min at 3,500 x g), frozen in liquid nitrogen, and stored at -80 °C.

All steps of purification were completed at 4 °C. The cell pellet was placed on ice and thawed for 30 min prior to resuspension in Buffer A: 50 mM sodium phosphate, pH 7.6, 300 mM NaCl, 2 mM imidazole, 5% (v/v) glycerol and 1 mM tris(2-carboxyethyl)phosphine (TCEP). For lysis, Buffer A was supplemented with 5 U mL^-1^ DNase, 5 U mL^-1^ RNase A and 0.2 mg mL^-1^ lysozyme (hen egg white, Sigma), 1.5 mM MgCl2, and EDTA-free protease inhibitor tablet (Roche), 250 µM phenylmethylsulfonyl fluoride (PMSF). Cells were homogenized with a French Press at 14,000 psi for 10 minutes inside a cold room operating at 4 °C. Cell debris was separated by centrifugation at 4 °C (25,000 x g, 30 min), and the supernatant was loaded onto a Talon cobalt affinity column (5 mL bed volume) equilibrated in Buffer A. The column was washed with the equilibration buffer for 10 column volumes, and protein was eluted with Buffer A with 500 mM imidazole. Protein-containing fractions were pooled and buffer exchanged into Buffer A via a centrifugal concentrator (30 kDa Amicon). Smt3-protease was added of a 350:1 mole ratio of tagged protein:Smt3 protease. Detagged protein was collected via the flowthrough of a second cobalt affinity column equilibrated with Buffer A. Further purification was performed with size exclusion chromatography on a HiLoad Superdex 200 pg preparative 16/600 column in Buffer A. All protein concentrations are given as monomer concentrations. All biophysical studies were performed in the following assay buffer except where noted in the cryo-EM sample preparation: 50 mM HEPES pH 7.6, 300 mM NaCl, 1 mM TCEP and 5% (v/v) glycerol.

### Small-angle X-ray scattering

X-ray scattering experiments were performed at the Cornell High Energy Synchrotron Source (CHESS) ID7A1 station. Data were collected using a 250 µm x 200 µm X-ray beam with an energy of 9.9 keV and a flux of ∼10^12^ photons s^-1^ mm^-1^ at the sample position. Small-angle X-ray scattering (SAXS) images were collected on an Eiger 4M detector covering a range of *q* = 0.009–0.55 Å^−1^. Here, the momentum transfer variable is defined as *q*=4π/*λ* sin*θ*, where *λ* is the X-ray wavelength and 2*θ* is the scattering angle. Data processing at the beamline was performed in BioXTAS RAW (Hopkins et al., 2017). Briefly, scattering images were integrated about the beam center and normalized by transmitted intensities measured on a photodiode beamstop. The integrated protein scattering profile, *I*(*q*), was produced by subtraction of background buffer scattering from the protein solution scattering. Radii of gyration (*Rg*) were estimated with Guinier analysis, and pair distance distribution analysis was performed with Bayesian indirect Fourier transformation (BIFT) (Hansen, 2000). Error bars associated with *Rg* values are curve-fitting uncertainties from Guinier analysis. Subsequent analysis was performed in MATLAB and utilized REGALS and other established protocols (Meisburger et al., 2021, 2016; Skou et al., 2014) The scattering of *Synechococcus* phage S-CBP4 α subunit was first measured over the concentration range of 4 – 40 µM. 8 µM was chosen for subsequent titration of nucleotides to maintain near physiological concentration (Ando et al., 2011; Parker, 2017). All nucleotide stocks were prepared with 15 mM MgCl2. For all titration experiments, background subtraction was performed with carefully matched buffer solutions containing identical concentrations of nucleotides following established protocols (Skou et al., 2014). For each measurement, 40 μL of sample were prepared fresh and centrifuged at 14,000×g at 4 °C for 10 min immediately before loading into an in-vacuum flow cell kept at 4 °C. For each protein and buffer solution, 20 × 2 s exposures were taken with sample oscillation to limit radiation damage then averaged together to improve signal. SVD was performed in MATLAB.

Size-exclusion chromatography-coupled SAXS (SEC–SAXS) experiments were performed using a Superdex 200 Increase 10/300 GL (24 mL) column operated by a GE Akta Purifier at 4 °C with the elution flowing directly into an in-vacuum X-ray sample cell. To account for a ∼10-fold dilution of the sample during elution, 40 μL sample was prepared at 80 μM protein in assay buffer with 200 µM TTP and 200 µM GDP for substrate. Sample was then centrifuged at 14,000×g for 10 min at 4 °C before loading onto a column pre-equilibrated in a matched buffer containing 200 µM TTP, 200 µM GDP. Samples were eluted at flow rates of 0.05 mL min^−1^ and 2-s exposures were collected throughout elution until the elution profile had returned to buffer baseline. Normalized, integrated scattering profiles were binned 6-fold in frame number and 4-fold in *q*, and scattering profiles of the elution buffer were averaged to produce a background-subtracted SEC–SAXS dataset. Singular value decomposition (SVD) was performed to determine the number of significant components, and the SEC-SAXS dataset was decomposed in the MATLAB implementation of REGALS (Meisburger et al., 2021).

Structural modeling was performed using the ATSAS package (Manalastas-Cantos et al., 2021) and AllosModFoXS (Schneidman-Duhovny et al., 2010; Weinkam et al., 2012) following previously established protocols (Ando et al., 2016, 2011; Meisburger et al., 2016; Thomas et al., 2019). Theoretical scattering curves of the cryo-EM model (Methods described below, PDB: 7urg) were calculated in CRYSOL (Svergun et al., 1995) with 50 spherical harmonics, 256 points between 0 and 0.5 Å−1, and the default electron density of water. The overall scale factor and solvation parameters were determined by fitting to the protein scattering curve extracted by REGALS. Disordered and missing residues were modeled in AllosModFoXS (Konarev et al., 2003; Manalastas-Cantos et al., 2021) with sampling of static structures consistent with the starting model.

### Cryo-EM grid preparation and data acquisition

QuantiFoil holey carbon R 1.2/1.3 300-mesh grids were glow discharged on a PELCO easiGlow system for 45 s with 15 mA current. Grid freezing was then performed on a FEI Vitrobot Mark IV with the chamber humidity set to 100% and the temperature set to 4 °C. 3 μL of sample (4 µM *Synechococcus* phage S-CBP4 α subunit in 50 mM HEPES pH 7.6, 150 mM NaCl, 7.55 mM MgCl2, 200 µM TTP, 200 µM GDP, 1 mM TCEP, 1% v/v glycerol) was applied onto the grid. The sample was blotted for 4 s and then immediately plunged into liquid ethane cooled by liquid nitrogen.

Data collection was performed at the Cornell Center for Materials Research (CCMR) on a Talos Arctica (Thermo Fisher Scientific) operating at 200 keV with a Gatan K3 direct electron detector and BioQuantum energy filter at a nominal magnification of 79,000 x (1.07 Å/pixel). A total of 856 movies was collected with a nominal defocus range from -0.6 μm to -2.0 μm and a total dose of 50 e^-^/Å^2^ over 50 frames (2.164 s total exposure time, 0.0435 s frame time, 26.96 e^-^/Å^2^/s dose rate).

### Cryo-EM data processing

Initial processing was performed in cryoSPARC v3.3.1 (Punjani et al., 2017). Patch motion correction and patch CTF estimation were performed on 856 movies. The resulting micrographs were manually curated based on statistics and visual inspection, and 432 micrographs were retained. 46 high quality micrographs were then selected, from which the blob picker routine was used to pick particles. The resulting 99k particles were extracted and subjected to 2D classification, and the top four unique 2D classes were selected and used as templates for template picking on the entire dataset. Due to the large variance in ice conditions in many of our micrographs, masks were manually defined for every micrograph, and particle picks outside the ideal ice region were excluded. The resulting 582k particles were extracted with a box size of 256 pixels binned to 128 pixels. Two rounds of 2D classification were performed, and only particles corresponding to the top 2D classes with secondary structure features were kept. The 203k remaining particles were re-extracted with 256-pixel box size and subjected to *ab initio* reconstruction and heterogeneous refinement into two classes. The top class containing 117k particles was then subjected to homogeneous refinement with C2 symmetry, which yielded a 4.04 Å map. Duplicate particles were removed using a minimum separation distance of 80 Å, and the remaining particles (108k) were subjected to a non-uniform refinement (Punjani et al., 2020) with C2 symmetry imposed and per-particle defocus and CTF parameter optimization enabled, which yielded a 3.57 Å map.

To employ Bayesian polishing in RELION-3 (Zivanov et al., 2018), the same 432 micrographs were motion-corrected in RELION 3.1 using its own implementation of MotionCorr2 (Zheng et al., 2017) with 5 by 5 patches. The particle .cs file from the cryoSPARC non-uniform refinement job was converted to star format using pyem (Asarnow, D., Palovcak, E., & Cheng, Y., 2019) with the micrograph path modified to that of the RELION motion-corrected micrographs. Particles were then re-extracted from RELION motion-corrected micrographs with a box size of 256 pixels. Due to the failure of RELION 3D auto-refine to converge on a reasonable structure from these particles, particles were imported back into cryoSPARC and subjected to homogeneous refinement. The particle .cs file from this refinement job was again converted to a star file using pyem. Relion_reconstruct was employed to reconstruct two half maps using the offset and angle information refined in cryoSPARC with the same half data split, and post-processing was performed on the resulting half maps using the refinement mask from cryoSPARC. Bayesian polishing (Zivanov et al., 2019) training and polishing jobs were then performed with this post-processing job as input. The resulting shiny particles were successfully refined to a 3.85 Å map with 3D auto-refine in RELION. After one round of CTF refinement (Zivanov et al., 2020) the particles were subjected to another round of Bayesian polishing, and the resulting shiny particles were imported back into cryoSPARC. Non-uniform refinement on shiny particles with C2 symmetry imposed and per-particle defocus and CTF parameter optimization enabled yielded the final 3.46-Å resolution map used for model building and analysis.

### AlphaFold prediction and atomic model building

The sequence for the *Synechococcus* phage S-CBP4 α subunit was retrieved from UniProt (Bateman et al., 2021) with accession number M1PRZ0. The sequence was used as input for AlphaFold2 prediction (Jumper et al., 2021) with the five default model parameters and a template date cutoff of 2020-05-14. As the five models were largely identical in the core region and differing only in the location of the C-terminal tail, the structure predicted with the first model parameter was used in the subsequent process. Similar procedures were used to predict structures of class Ø ferritin-like proteins, ATP-cone-containing α sequences, and class II and III α sequences described in the text.

The predicted structure of the *Synechococcus* phage S-CBP4 α subunit was first processed and docked into the unsharpened cryo-EM map in phenix. The 25 N-terminal residues and 45 C-terminal residues were then manually removed due to lack of cryo-EM density, and residues 26-426 were retained in the model. We observed unmodeled density at the specificity site, and based on solution composition, we modeled a TTP molecule. A structure of TTP coordinating a magnesium ion was extracted from the crystal structure of *Bacillus subtilis* RNR (pdb: 6mt9) (Thomas et al., 2019) and rigid body fit into the unmodeled density in Coot (Emsley and Cowtan, 2004). The combined model was refined with unsharpened and sharpened maps using phenix.real_space_refine (Afonine et al., 2018; Liebschner et al., 2021), with a constraint applied on the magnesium ion coordinated by the triphosphate in TTP according to the original configuration. Residue and loop conformations in the resulting structure were manually adjusted in Coot to maximize fit to map and input for an additional round of real-space refinement in phenix with an additional restraint for the disulfide bond between C30 and C196. The atomic coordinates and maps have been deposited to the Protein Data Bank and EM Data Bank under accession codes 7urg and EMD-26712. Due to the weak density for the magnesium ion, it was removed from the atomic coordinates when deposited into PDB.

### EM Data collection, processing, and refinement

| Data collection and processing |  |
| --- | --- |
| Microscope | Talos Arctica |
| Camera | K3 |
| Magnification | 79,000 |
| Voltage (keV) | 200 |
| Electron exposure ($e^-/\text{\AA}^2$) | 50 |
| Defocus range ($\mu\text{m}$) | -0.6 to -2.0 |
| Pixel size ($\text{\AA}$) | 1.07 |
| Micrographs used (no.) | 432 |
| Initial particles (no.) | 581,884 |
| Final particles (no.) | 107,885 |
| Symmetry imposed | C2 |
| Map resolution (Å)<br>FSC threshold | 3.46<br>(0.143) |
| Map resolution range (Å) | 3.0 – 7.0 (75%) |
| **Refinement** |  |
| Initial model used | AlphaFold2 prediction for UniProt entry M1PRZ0 |
| Model resolution (Å)<br>FSC threshold | 3.6<br>(0.5) |
| Map sharpening B factor (Å<sup>2</sup>) | -171 |
| Model composition<br>Non-hydrogen atoms<br>Protein residues<br>Ligands | 6260<br>802<br>TTP |
| B factors (Å<sup>2</sup>)<br>Protein<br>Ligand | 74.37<br>67.23 |
| R.m.s. deviations<br>Bond lengths (Å)<br>Bond angles (°) | 0.004<br>0.937 |
| Validation<br>MolProbity score<br>Clashscore<br>Poor rotamers (%) | 1.47<br>4.69<br>0.00 |
| Ramachandran plot<br>Favored (%)<br>Allowed (%)<br>Disallowed (%) | 96.49<br>3.51<br>0.00 |

## Data Availability

The cryo-EM map has been deposited in the Electron Microscopy Data Bank under accession code EMD- 26712, and the model has been deposited in the Protein Data Bank under accession code 7urg.

## Author Contributions

NA and CJ conceived of the project, designed and supervised research, and analyzed data. AAB performed bioinformatic analyses, purified and expression protein, performed SAXS analysis, and analyzed and curated all project results. MS devised bioinformatic workflow, performed computational experiments, and assisted in data analysis and manuscript preparation. DX performed bioinformatic analyses and cryo-EM data acquisition and analysis. AAB, DX, and NA wrote the paper with contributions from MS and CJ. All authors edited the manuscript.

## Acknowledgments

The authors are grateful to Drs. Will Thomas and Steve Meisburger for helpful discussions and Drs. Richard Gillilan and Qingqiu Huang for assistance at the 1D7A beamline at CHESS. SAXS was conducted at the Center for High Energy X-ray Sciences (CHEXS), which is supported by the National Science Foundation (NSF) under award DMR-1829070, and the Macromolecular Diffraction at CHESS (MacCHESS) facility, which is supported by award 1-P30-GM124166-01A1 from the National Institute of General Medical Sciences (NIGMS), National Institutes of Health (NIH), and by New York State’s Empire State Development Corporation (NYSTAR). Cryo-EM work was done using the Cornell Center for Materials Research (CCMR) Shared Facilities, which are supported through the NSF MRSEC program (DMR-1719875). This project was undertaken with the assistance of resources and services from the National Computational Infrastructure (NCI), which is supported by the Australian Government, as well as with the BioHPC resource at the Cornell Institute of Biotechnology. We acknowledge the ARC Centre of Excellence for Innovations in Peptide and Protein Science (CE200100012), the ARC Centre of Excellence in Synthetic Biology (CE200100029). This work was supported by NSF grant MCB-1942668 (to N.A.) and startup funds from Cornell University (to N.A.).

---

**Figure 2 – figure supplement 1.**
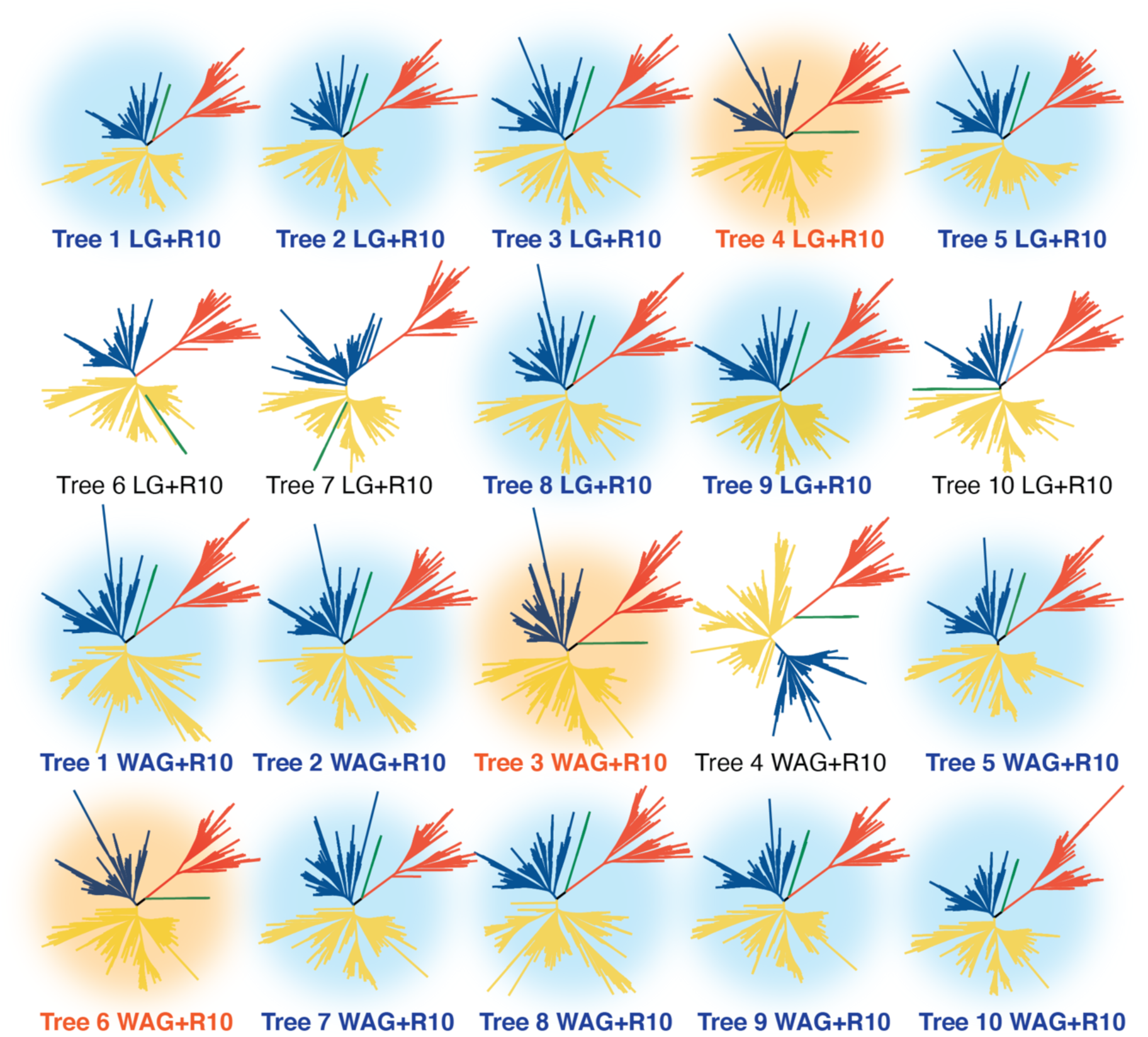
All 20 inferred phylogenies of the ribonucleotide reductase family shown in the unrooted representation. Green, red, yellow, and blue branches are clade Ø and classes III, II, and I, respectively. Trees are labeled with the respective matrices of evolution used for the phylogenetic inference. The tree shown in Figure 2 and referenced throughout this work is tree 1. Trees with the light blue background support the overall topology shown in Figure 2 of the major classes. Trees with orange background show the second most frequently observed topology, where the midpoint places the class Ø clade ancestral to the class III clade. The four trees without a colored background cannot be categorized into either of the previously described topologies.

**Figure 4 – figure supplement 1.**
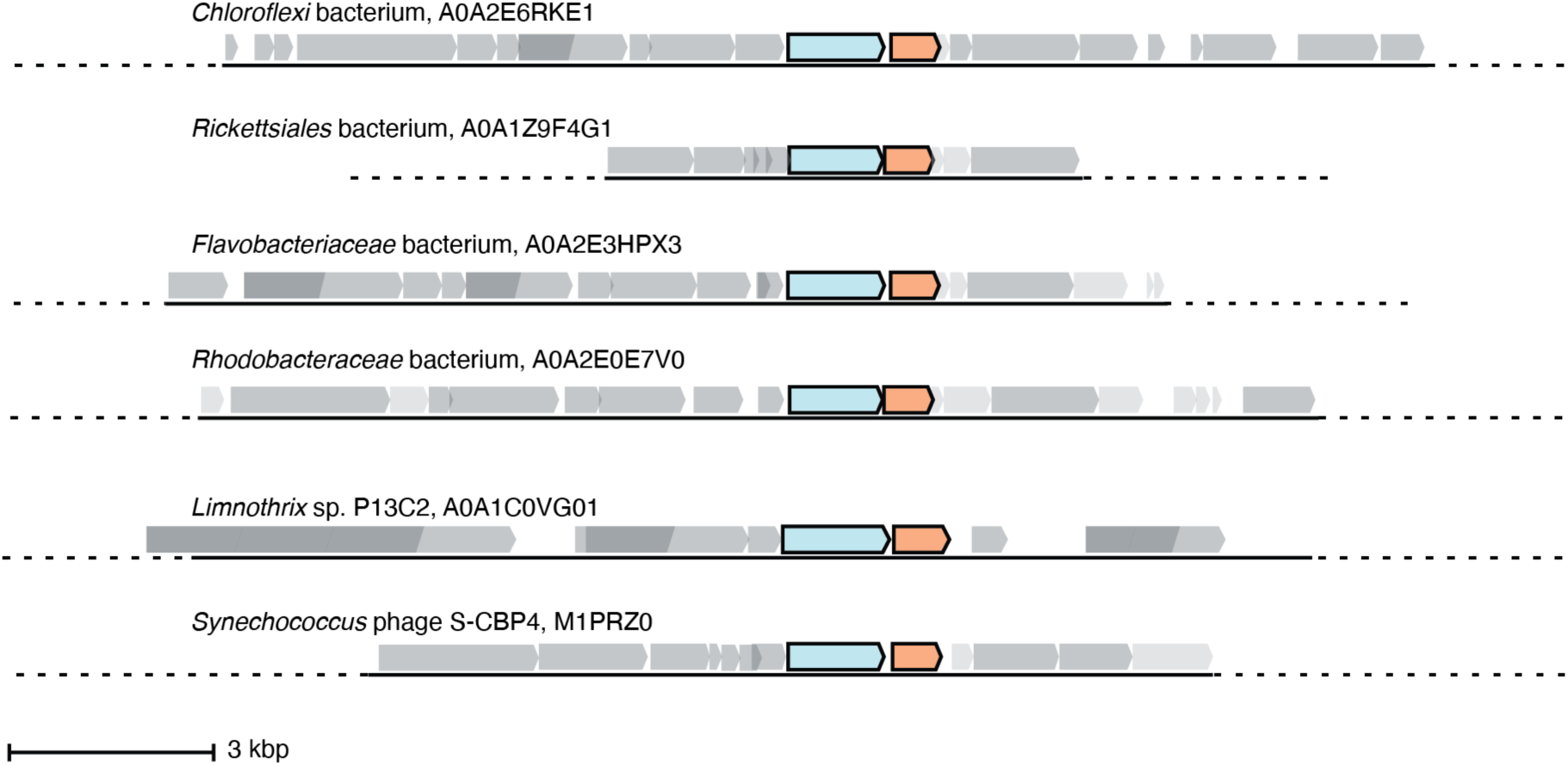
Six representative operons of class Ø sequences. In sequenced organisms with high genome coverage, the α gene (cyan) is followed by a gene for a small ferritin-like protein (orange). Of the 35 sequences in the class Ø clade, 24 are from sequenced genomes with greater coverage than a contig. 18 of these have annotated ferritin-like sequences (IPR009078) directly downstream of the α gene in the operon, and 6 have unannotated genes that share homology with the class Ø ferritin-like gene directly downstream of the α gene.

**Figure 4 – figure supplement 2.**
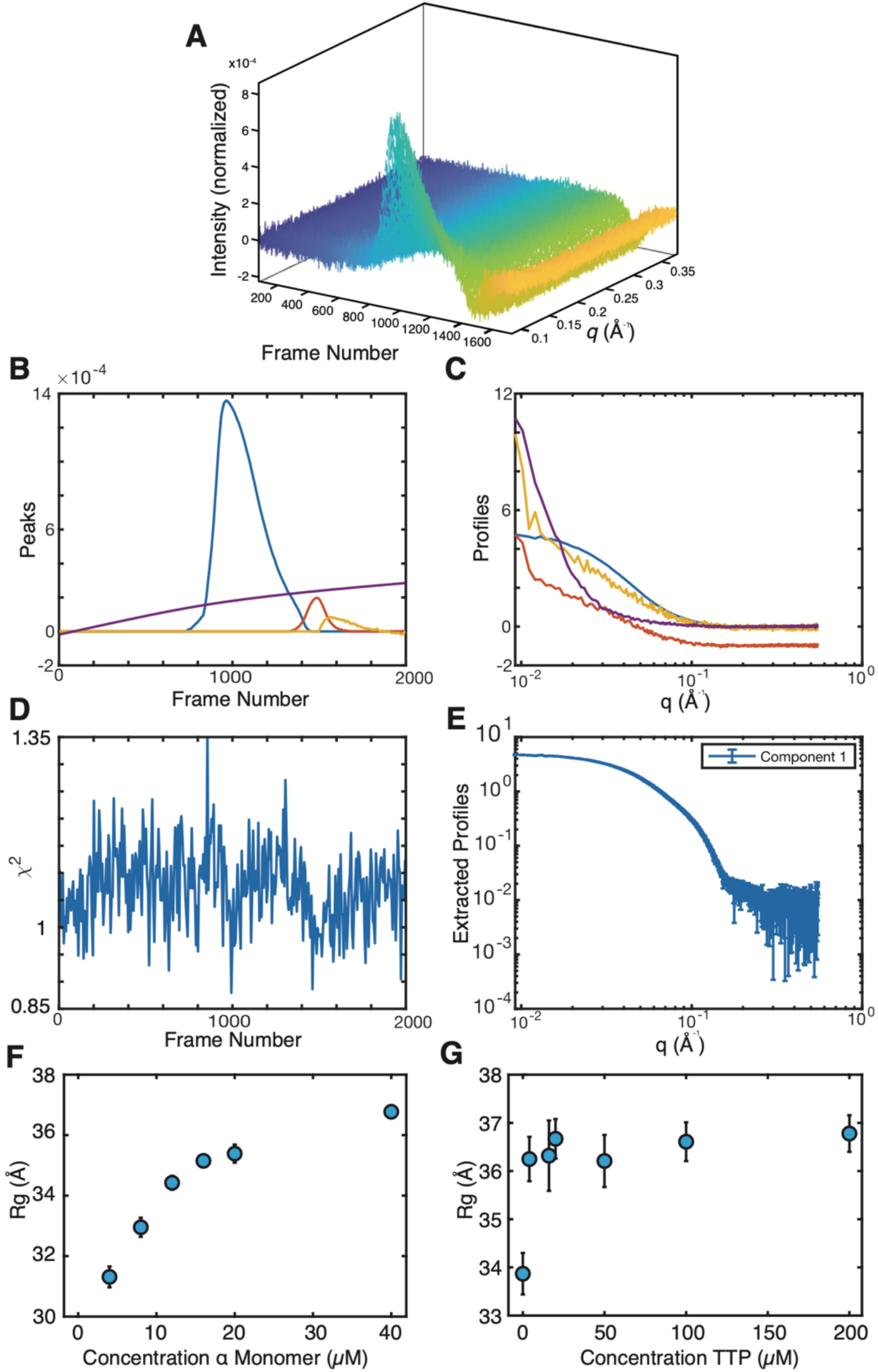
*Synechococcus* phage S-CBP4 α is a stable dimer in the presence of nucleotides. (**A**) Size exclusion chromatography-coupled SAXS (SEC-SAXS) data collected on the *Synechococcus* phage S-CBP4 α subunit in the presence of 200 µM TTP and 200 µM GDP. Singular value decomposition (SVD) of the background-subtracted SEC-SAXS data contained 4 significant singular values. REGALS (Meisburger et al., 2021) was used to decompose the dataset into (**B**) elution profiles and (**C**) SAXS profiles for a sloping background component (purple), a single protein component (blue), and 2 trailing components (red and yellow) representing buffer mismatch and radiation damage. (**D**) Residuals of REGALS calculation from SEC-SAXS dataset. (**E**) Extracted SAXS profile for the protein component. (**F**) The radius of gyration (*Rg*) of *Synechococcus* phage S-CBP4 α in the absence of nucleotides as a function of protein concentration in solution shows a monomer-dimer equilibrium. (**G**) The *Rg* of 8 µM *Synechococcus* phage S-CBP4 α in the presence of 200 µM GDP as a function of TTP concentration in solution is consistent with a stable dimer. Error bars represent uncertainties from Guinier analysis, and where not visible are smaller than the data marker. The theoretical *Rg* values of *Synechococcus* phage S-CBP4 α (based on the cryo-EM model with N- and C-termini added) are 26 Å and 37 Å, respectively, for monomer and dimer.

**Figure 4 – figure supplement 3.**
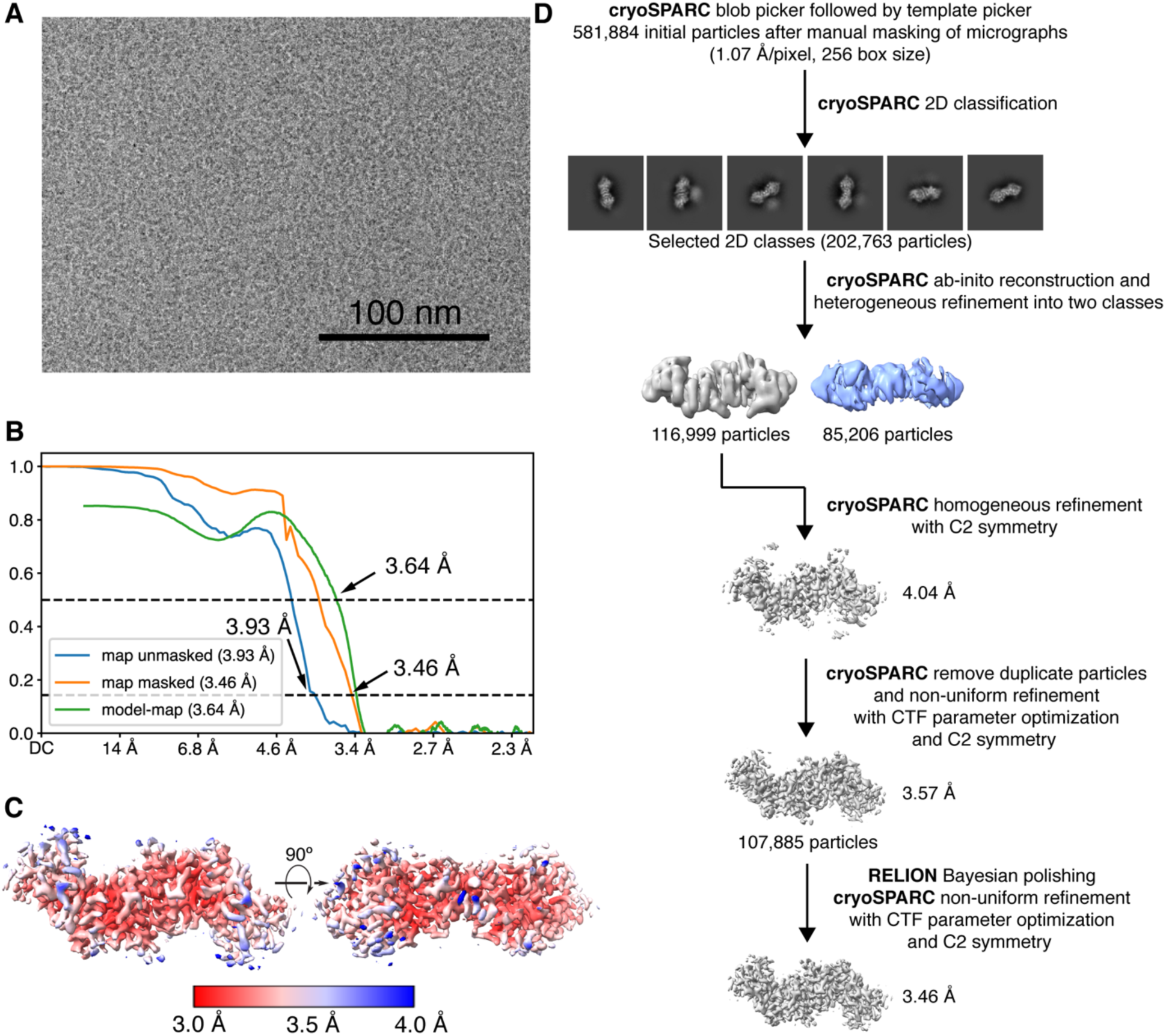
Cryo-EM image processing workflow for reconstructing the 102 kDa *Synechococcus* phage S-CBP4 α2 dimer. (**A**) Representative micrograph used in processing. (**B**) Fourier shell correlation (FSC) curves for the final map with and without mask reported by cryoSPARC shown as blue and orange curves. The model-map FSC curve (green) was calculated for the deposited model (PDB: 7urg) and the sharpened map using PHENIX. (**C**) Local resolution estimation computed in cryoSPARC for the final map. (**D**) Data processing scheme used to obtain the final map.

**Figure 4 – figure supplement 4.**
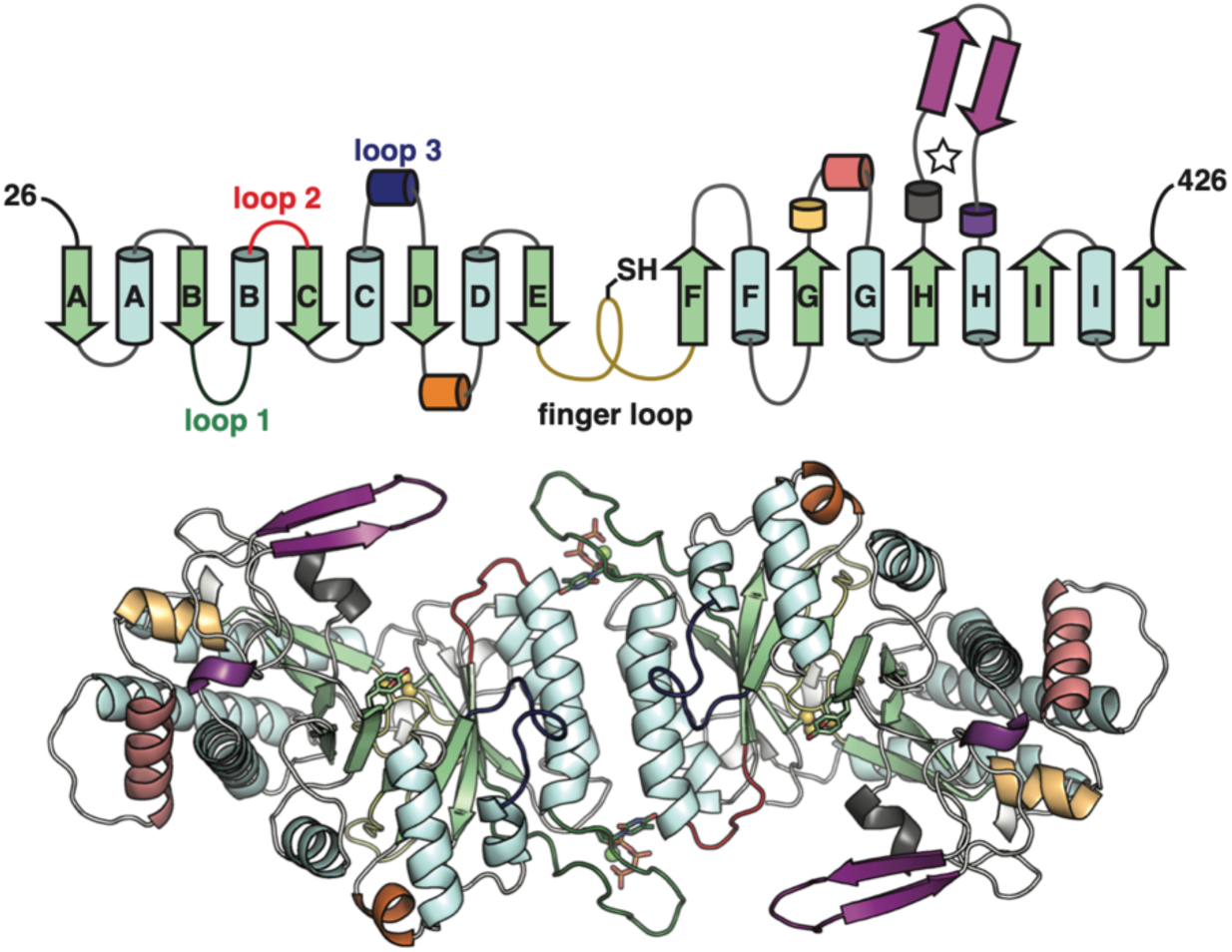
The class Ø α subunit features the most minimal RNR architecture discovered to-date. Top: Topology of the *Synechococcus* phage S-CBP4 α subunit monomer based on the cryo-EM model the minimal components of the RNR catalytic barrel colored as in Figure 1A. Only a few insertions are observed in the class Ø α subunit (yellow orange, orange, and violet secondary structures). Bottom: The cryo-EM model of the *Synechococcus* phage S-CBP4 α dimer with the same coloring as above.

**Figure 4 – figure supplement 5.**
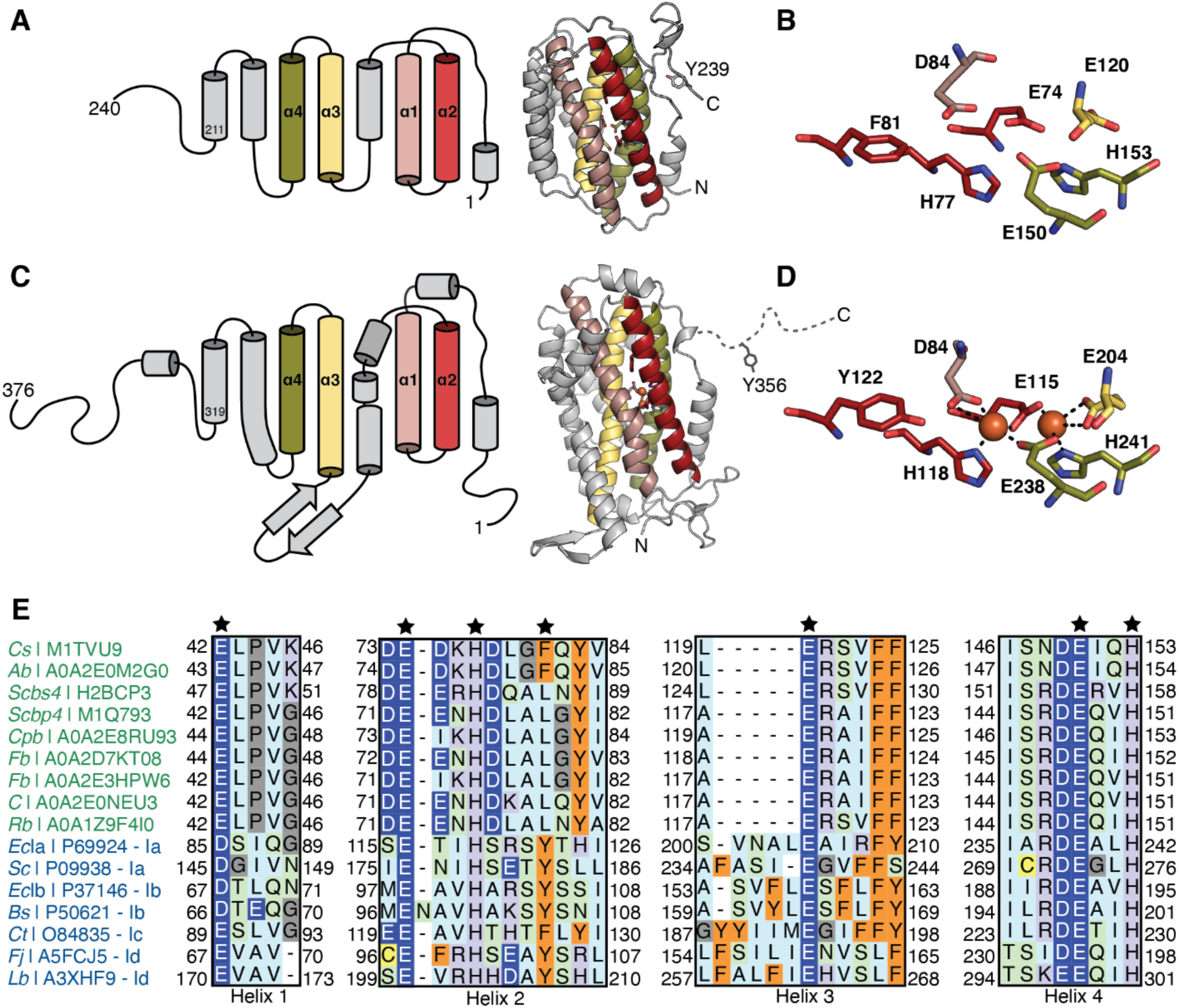
Comparison of class Ø ferritin-like proteins with class I RNR β subunits. The four-helical bundle of the ferritin fold is colored in red, pink, yellow, and olive. (**A**) Topology (left) of an AlphaFold2 model (right) of a representative class Ø ferritin-like protein (Unitprot: M1TVU9) and (**B**) a close-up of the residues that would putatively form the metal-binding motif. In this sequence, a redoxinert phenylalanine (F81) is found in the *i*+4 position from first histidine in the motif (H77). (**C**) Topology (left) of the crystal structure (right) of the *E. coli* class Ia RNR β subunit (PDB: 1piy) and (**D**) a close-up of the residues that form the diiron center. The site of the radical is a tyrosine (Y122) in the *i*+4 position from the first histidine in the motif (H118). (**E**) Sequence alignment of representative class Ø ferritin-like protein and class I RNR β subunits. Class Ø colored in green: *Cs,* Cyanophage SS120-1; *Ab, Alphaproteobacteria bacterium; Scbs4, Synechococcus* phage *S-CBS4; Scbp4, Synechococcus* phage *S-CBP4; Cpb, Candidatus Poribacteria bacterium; Fb, Flavobacteriales bacterium; Fb, Flavobacteriaceae bacterium; C, Coraliomargarita sp.; Rb, Rickettsiales bacterium* TMED131. Class I colored in blue: Class Ia: *EcIa, Escherichia coli* (strain K12)*; Sc, Saccharomyces cerevisiae* (strain ATCC 204508 / S288c). Class Ib: *EcIb, Escherichia coli* (strain K12)*; Bs, Bacillus subtilis* (strain 168). Class Ic *Ct Chlamydia trachomatis* (strain D/UW-3/Cx). Class Id: *Fj Flavobacterium johnsoniae* (strain ATCC 17061); *Lb, Leeuwenhoekiella blandensis* (strain CECT 7118). A conserved E/D and EXXH metal-binding motif is observed in each helical pair (1 + 2; 3 + 4). The *i*+4 position from the histidine in helix 2 is Phe or Leu in class Ø ferritin-like proteins, much like in class Ic β subunits.

**Figure 5 – figure supplement 1.**
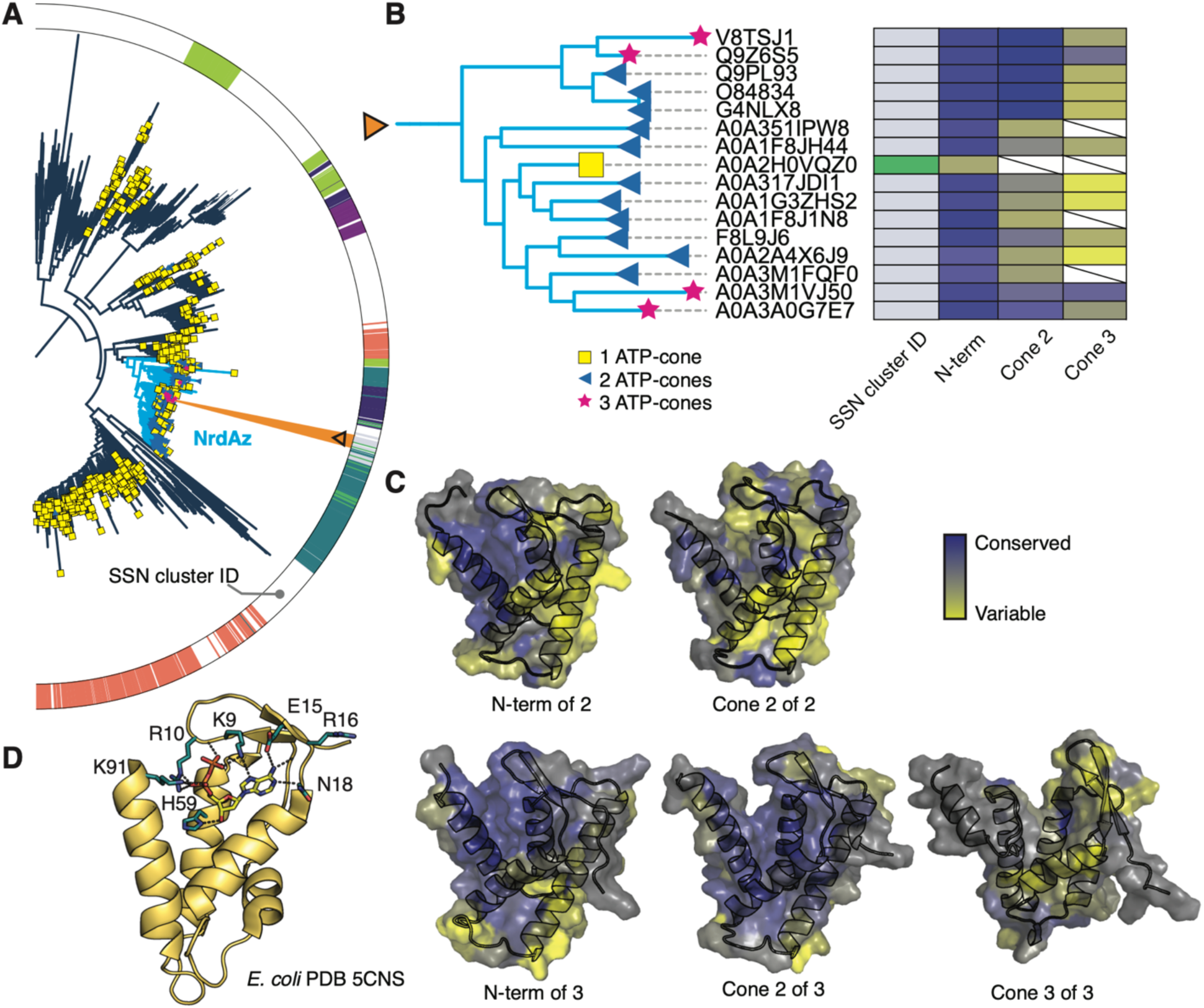
Inner copies in tandem ATP-cones diverge from the canonical motif while N-terminal cones maintain homology. (**A**) The class I phylogeny pruned from Figure 2. Tips denote number of ATP-cones in the sequence. The color strip corresponds to the SSN cluster ID as shown in Figure 5. The light blue branches correspond to NrdAz sequences. The sequences in the orange wedge refers are shown in panel B. (**B**) An example of a tandem ATP-cone containing clade pruned from the class I phylogeny (orange wedge in panel A). The colors in the leftmost color strip represents the SSN cluster ID from Figure 5. The second, third, and fourth color strips show heatmaps of the individual bit score of each copy of the tandem ATP-cones when aligned to the Pfam03477 HMM. (**C**) AlphaFold2 models of ATP-cone domains from the double- and triple-domain containing sequences UniProt IDs A0A2A4X6J9 A0A1F8J1N8 (top set, *Chlamydiae* bacterium) and (bottom set, *Candidatus* Aerophobetes bacterium). Sequence conservation was calculated separately for all sequences in our dataset with two or three ATP-cones in Consurf (Landau et al., 2005). Conservation levels were then mapped onto the AlphaFold2 models, where residues of high conservation are purple and low conservation are yellow. (**D**) For comparison, the ATP-cone from *E. coli* class Ia α is shown with dATP bound (yellow). Key interacting residues are shown as teal sticks.

**Figure 6 – supplement 1.**
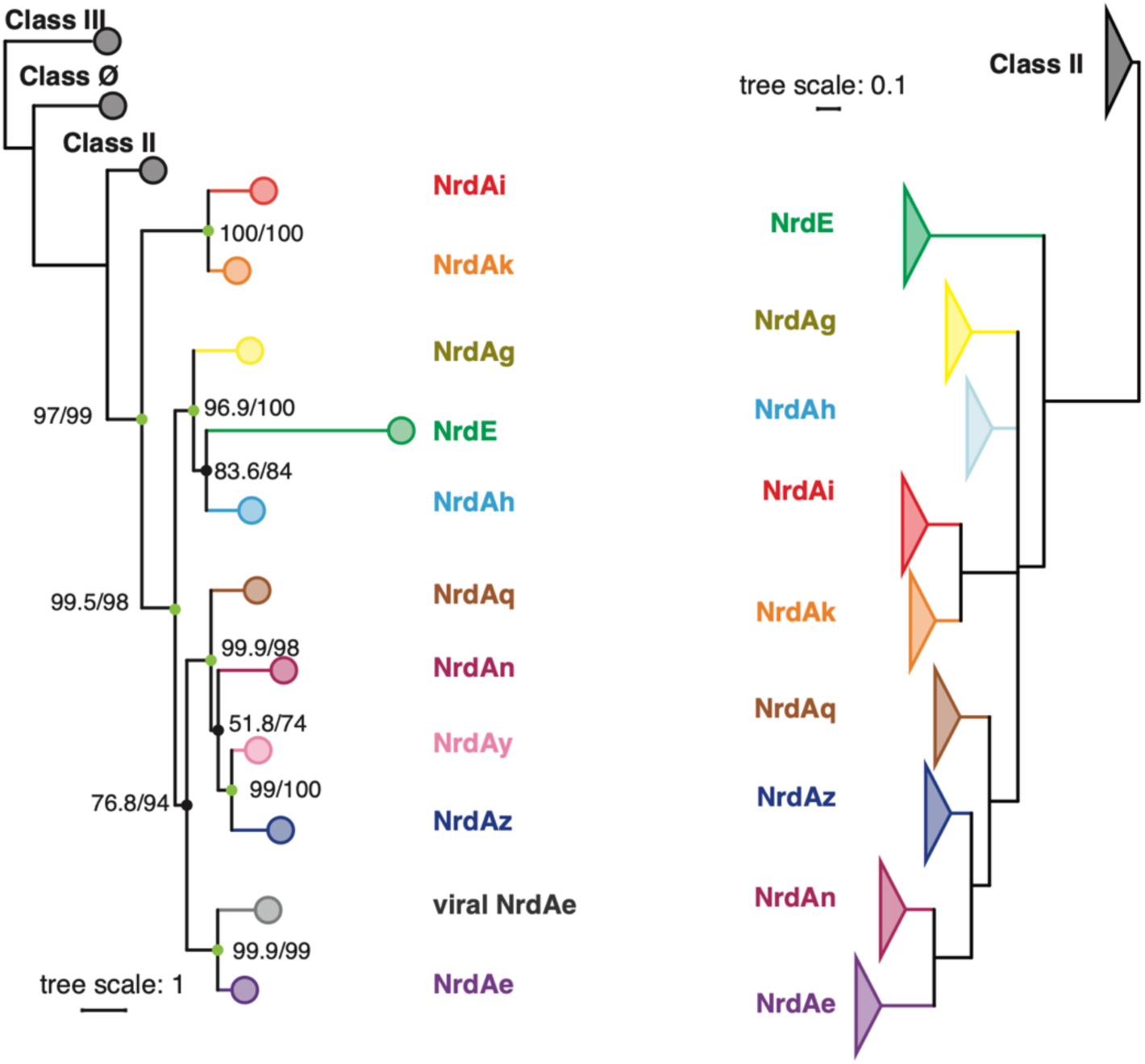
Comparison of class I RNR phylogenies. Left: the same as shown in Figure 6. Right: the tree published by (Martínez-Carranza et al., 2020) shown in collapsed form for direct comparison.

**Figure 6 – supplement 2.**
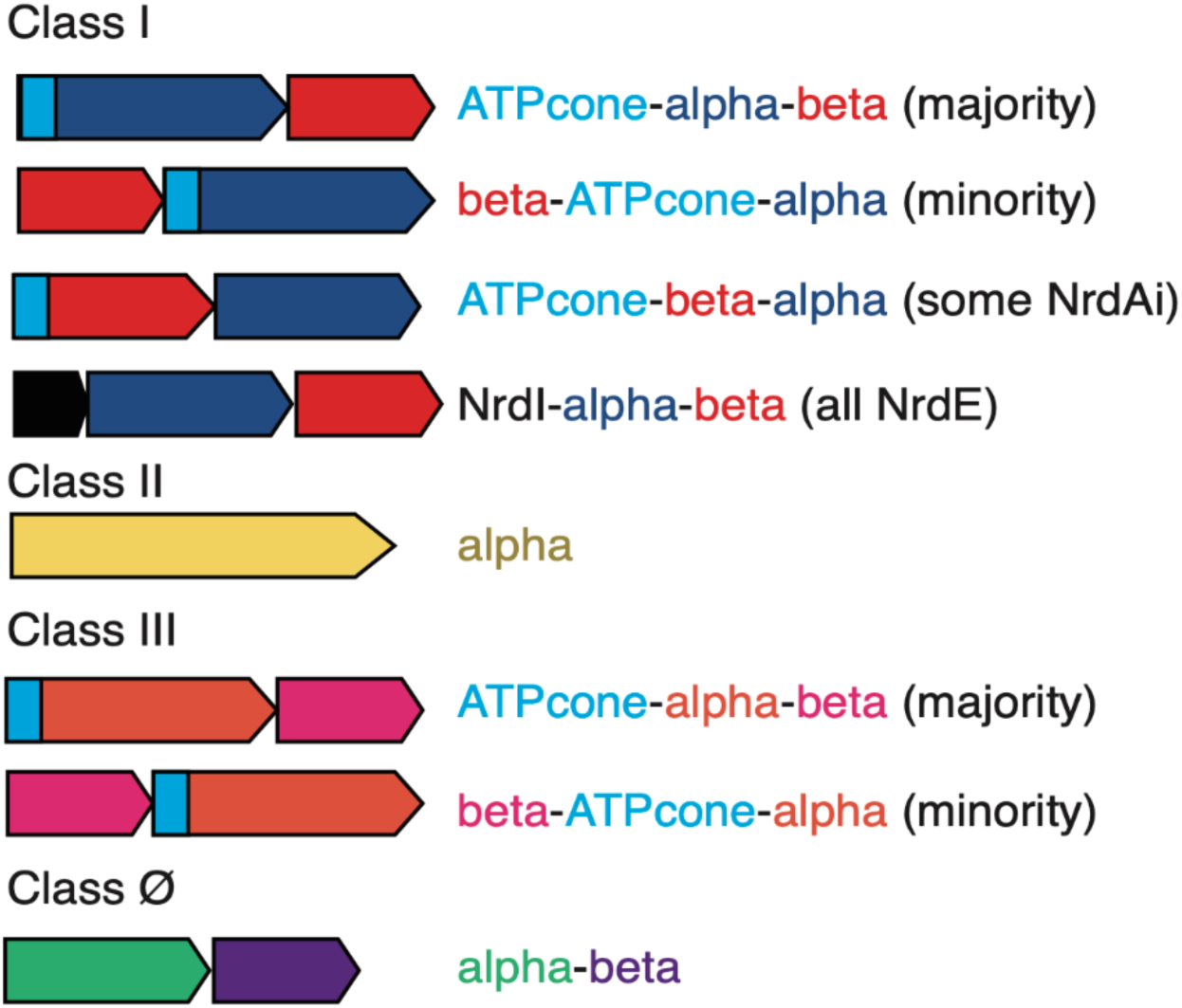
Overview of RNR operon organization. For sequences that contain ATP-cones, the location is shown in light blue.

**Figure 6 – supplement 3.**
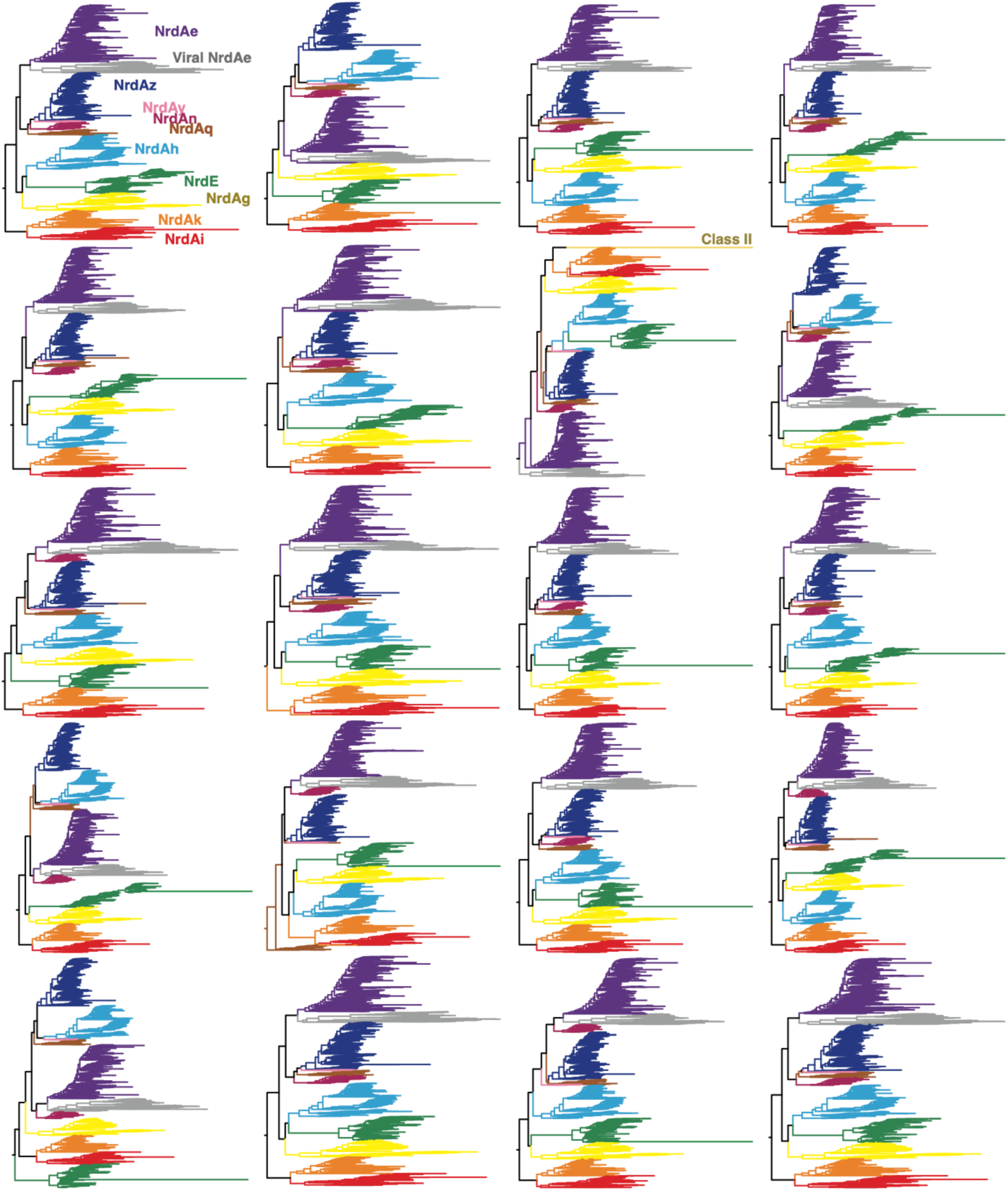
All 20 topologies of the class I clade. The first ten trees from left to right, top to bottom, were calculated with the LG+R10 model of evolution, and the latter ten trees were calculated with the WAG+R10 model of evolution. Tree 1, shown in Figure 2 and examined in Figure 6, contains the color codes for the phylogenetic subclasses of RNR.

**Figure 7 – figure supplement 1.**
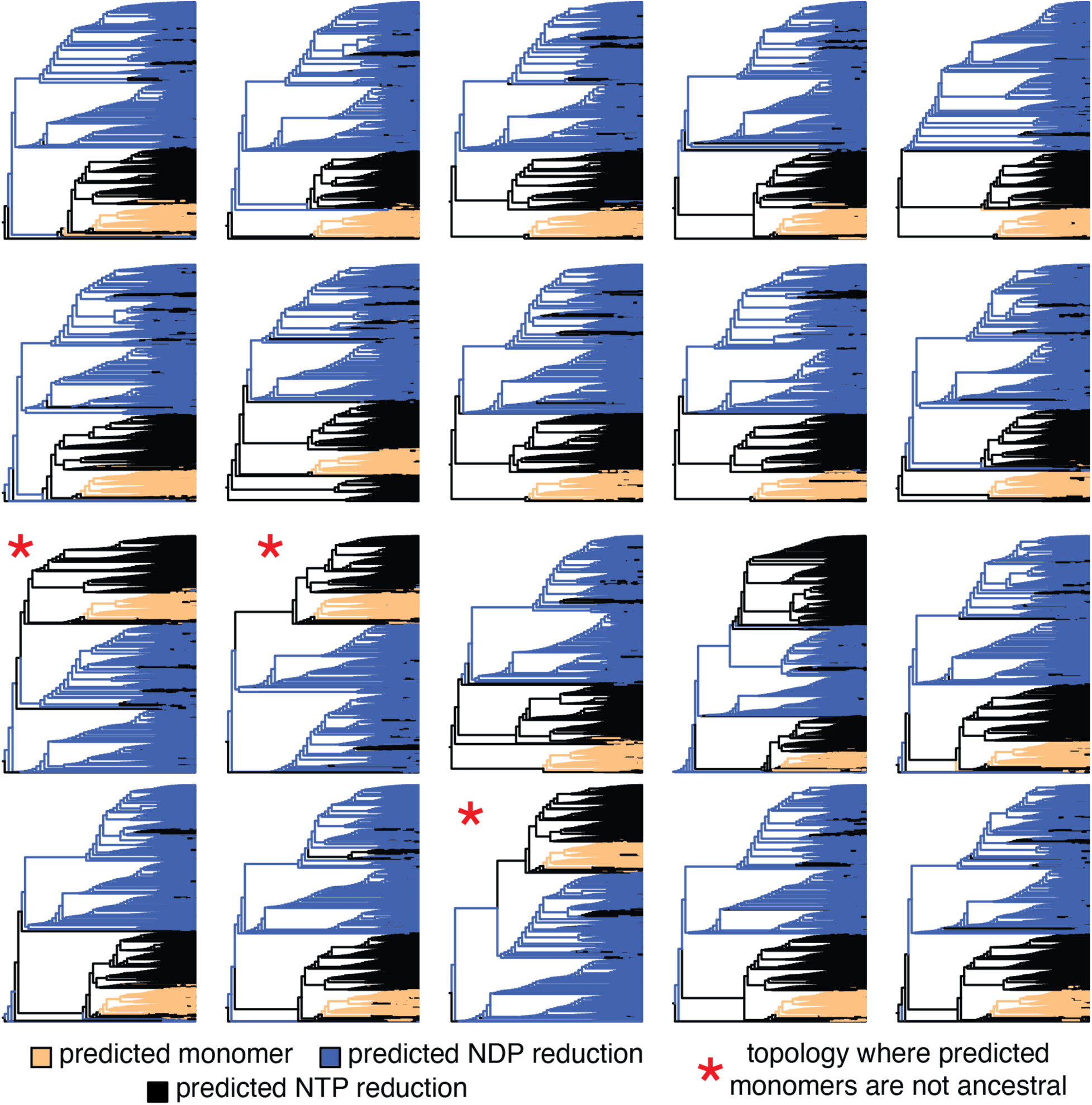
The topologies of the class II clade from all 20 tree inferences. The top ten trees were calculated under the LG+R10 model of evolution, and the bottom ten trees were calculated under the WAG+R10 model of evolution. The peach branches correspond to sequences that contain the monomer-conferring insertion between βB and αB and are in clades that have a representative sequence predicted by AlphaFold2 to be monomeric. Blue branches are sequences that are predicted to be specific for nucleotide diphosphate (NDP) reduction and black branches are sequences that are predicted to reduce nucleotide triphosphate substrates (NTPs). Note that all predicted monomer sequences are predicted to be NTP-specific. The three outlier topologies that do not predict the ancestral branch to the class II clade as NTP-specific monomer enzymes are annotated with a red asterisk. IscU-like domains (DX)

**Figure 7 – figure supplement 2.**
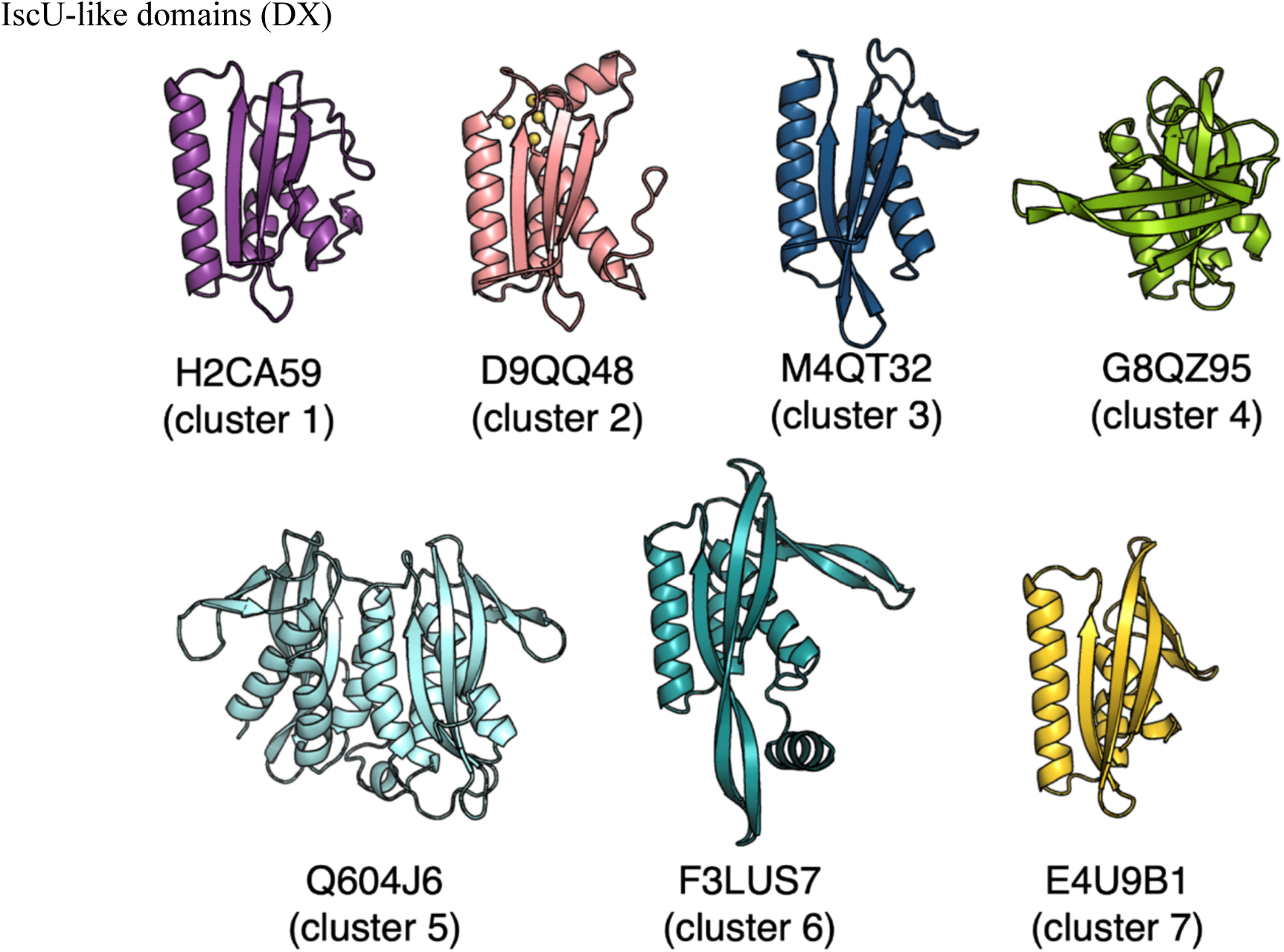
AlphaFold2 predictions of IscU-like C-terminal domains from representative sequences in each major cluster of the SSN. The UniProt IDs are shown for each sequence. In order of cluster number, the sequences correspond to the organisms *Leptonema illini* (H2CA59)*, Acetohalobium arabaticum* (D9QQ48)*, Loktanella* phage pCB2051-A (M4QT32)*, Owenweeksia hongkongensis* (G8QZ95)*, Methylococcus capsulatus* (Q604J6)*, Rubrivivax benzoatilyticus* JA2 (F3LUS7), and *Oceanithermus profundus* (E4U9B1), respectively.

**Figure 8 – figure supplement 1.**
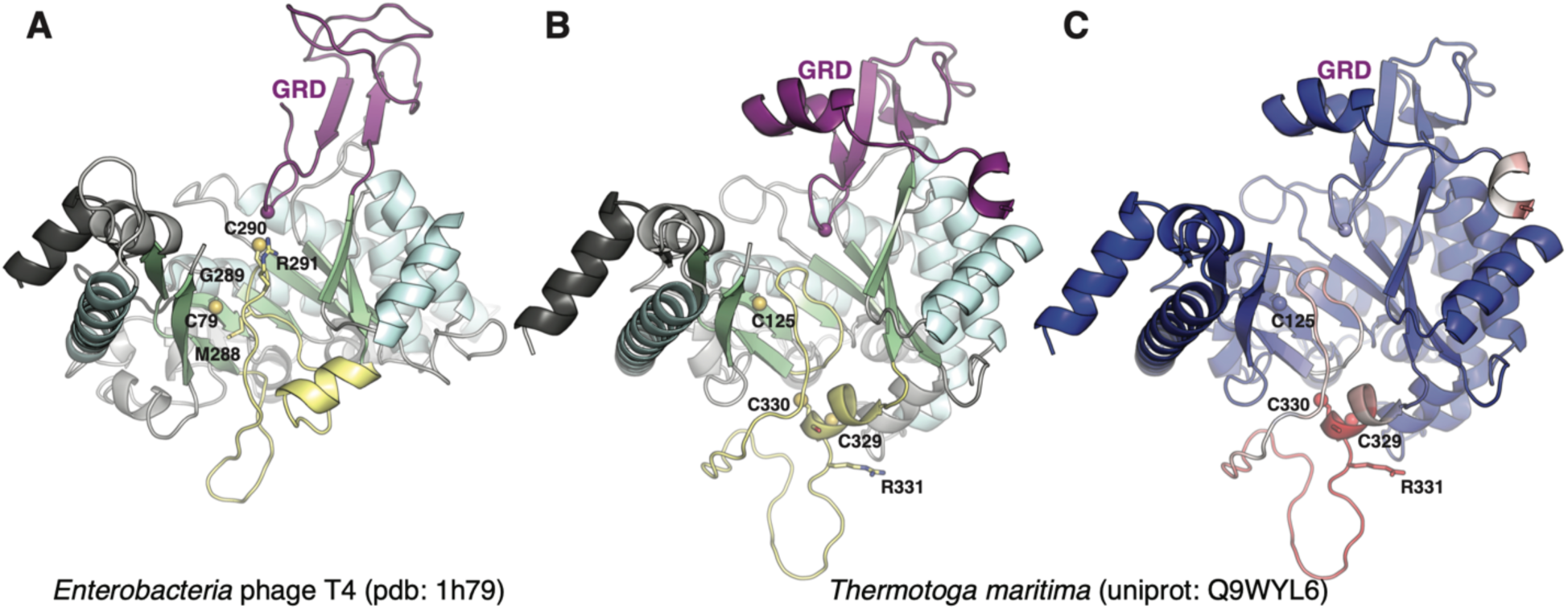
Class III sequences with crystal structures. The finger-loop is shown in yellow, and the cysteine sulfur atoms are shown as spheres. The glycyl radical domain (GRD) is shown in purple, and the alpha-carbon of the radical site is shown as a sphere. (**A**) Structure of class III RNR from *Enterobacteria* phage T4 (PDB 1h79). A single cysteine is at the tip of the finger loop. (**B**) AlphaFold2 prediction of *Thermotoga maritima* is identical to the known structure (PDB 4u3e) except with the missing region in the finger loop modeled. (**C**) Model in panel B colored by AlphaFold pLDDT confidence score where red is low confidence (pLDDT = 13.72) and blue is high (pLDDT = 100.96).

## Notes

### Competing Interest Statement

The authors have declared no competing interest.

## References

Andersson J, Westman M, Hofer A, Sjoberg BM. 2000. Allosteric regulation of the class III anaerobic ribonucleotide reductase from bacteriophage T4. J Biol Chem 275:19443–19448.

Ando N, Brignole EJ, Zimanyi CM, Funk MA, Yokoyama K, Asturias FJ, Stubbe J, Drennan CL. 2011. Structural interconversions modulate activity of Escherichia coli ribonucleotide reductase. Proceedings of the National Academy of Sciences 108:21046–21051.

Aravind L, Wolf YI, Koonin EV. 2000. The ATP-Cone: An evolutionarily mobile, ATP-binding regulatory domain. J Mol Microbiol Biotechnol 2:191–194.

Atkinson HJ, Morris JH, Ferrin TE, Babbitt PC. 2009. Using sequence similarity networks for visualization of relationships across diverse protein superfamilies. PLoS One 4:e4345.

Aurelius O, Johansson R, Bågenholm V, Lundin D, Tholander F, Balhuizen A, Beck T, Sahlin M, Sjöberg B-M, Mulliez E, Logan DT. 2015. The Crystal Structure of Thermotoga maritima Class III Ribonucleotide Reductase Lacks a Radical Cysteine Pre-Positioned in the Active Site. PLoS One 10:e0128199.

Ayala-Castro C, Saini A, Outten FW. 2008. Fe-S cluster assembly pathways in bacteria. Microbiol Mol Biol Rev 72:110–25, table of contents.

Belshaw R, Pybus OG, Rambaut A. 2007. The evolution of genome compression and genomic novelty in RNA viruses. Genome Res 17:1496–1504.

Bester AC, Roniger M, Oren YS, Im MM, Sarni D, Chaoat M, Bensimon A, Zamir G, Shewach DS, Kerem B. 2011. Nucleotide deficiency promotes genomic instability in early stages of cancer development. Cell 145:435–446.

Blaesi EJ, Palowitch GM, Hu K, Kim AJ, Rose HR, Alapati R, Lougee MG, Kim HJ, Taguchi AT, Tan KO, Laremore TN, Griffin RG, Krebs C, Matthews ML, Silakov A, Bollinger JM Jr, Allen BD, Boal AK. 2018. Metal-free class Ie ribonucleotide reductase from pathogens initiates catalysis with a tyrosine-derived dihydroxyphenylalanine radical. Proc Natl Acad Sci U S A 115:10022–10027.

Bollinger JM Jr, Jiang W, Green MT, Krebs C. 2008. The manganese(IV)/iron(III) cofactor of Chlamydia trachomatis ribonucleotide reductase: structure, assembly, radical initiation, and evolution. Curr Opin Struct Biol 18:650–657.

Booker S, Licht S, Broderick J, Stubbe J. 1994. Coenzyme B12-dependent ribonucleotide reductase: evidence for the participation of five cysteine residues in ribonucleotide reduction. Biochemistry 33:12676–12685.

Cotruvo JA, Stubbe J. 2011. Class I ribonucleotide reductases: metallocofactor assembly and repair in vitro and in vivo. Annu Rev Biochem 80:733–767.

Dridi N, Hadzagic M. 2019. Akaike and Bayesian Information Criteria for Hidden Markov Models. IEEE Signal Process Lett 26:302–306.

Dwivedi B, Xue B, Lundin D, Edwards RA, Breitbart M. 2013. A bioinformatic analysis of ribonucleotide reductase genes in phage genomes and metagenomes. BMC Evol Biol 13:33.

Eliasson R, Pontis E, Jordan A, Reichard P. 1999. Allosteric control of three B12-dependent (class II) ribonucleotide reductases. Implications for the evolution of ribonucleotide reduction. J Biol Chem 274:7182–7189.

Eliasson R, Pontis E, Jordan A, Reichard P. 1996. Allosteric regulation of the third ribonucleotide reductase (NrdEF enzyme) from enterobacteriaceae. J Biol Chem 271:26582–26587.

Fontecave M, Eliasson R, Reichard P. 1989. Oxygen-sensitive ribonucleoside triphosphate reductase is present in anaerobic Escherichia coli. Proc Natl Acad Sci U S A 86:2147–2151.

Funk MA. 2015. Structural Studies of Radical Enzymes in Bacterial Central Metabolism (PhD). MIT. Funk MA, Zimanyi CM, Andree GA, Hamilos AE, Drennan CL. 2021. fHow ATP and dATP act as molecular switches to regulate enzymatic activity in the prototypic bacterial class Ia ribonucleotide reductase. bioRxiv. doi:10.1101/2021.07.31.454598

Gao E-B, Huang Y, Ning D. 2016. Metabolic Genes within Cyanophage Genomes: Implications for Diversity and Evolution. Genes 7. doi:10.3390/genes7100080

Goldman AD, Kaçar B. 2021. Cofactors are Remnants of Life’s Origin and Early Evolution. J Mol Evol 89:127–133.

Greene BL, Kang G, Cui C, Bennati M, Nocera DG, Drennan CL, Stubbe J. 2020. Ribonucleotide Reductases: Structure, Chemistry, and Metabolism Suggest New Therapeutic Targets. Annu Rev Biochem 89:45–75.

Greene BL, Taguchi AT, Stubbe J, Nocera DG. 2017. Conformationally Dynamic Radical Transfer within Ribonucleotide Reductase. J Am Chem Soc 139:16657–16665.

Haghverdi L, Büttner M, Wolf FA, Buettner F, Theis FJ. 2016. Diffusion pseudotime robustly reconstructs lineage branching. Nat Methods 13:845–848.

Harrison AO, Moore RM, Polson SW, Wommack KE. 2019. Reannotation of the Ribonucleotide Reductase in a Cyanophage Reveals Life History Strategies Within the Virioplankton. Front Microbiol 10:134.

Hie BL, Yang KK, Kim PS. 2022. Evolutionary velocity with protein language models predicts evolutionary dynamics of diverse proteins. Cell Syst. doi:10.1016/j.cels.2022.01.003

Holmgren A, Sengupta R. 2010. The use of thiols by ribonucleotide reductase. Free Radical Biology and Medicine 49:1617–1628.

Jabłońska J, Tawfik DS. 2021. The evolution of oxygen-utilizing enzymes suggests early biosphere oxygenation. Nat Ecol Evol 5:442–448.

Johansson R, Jonna VR, Kumar R, Nayeri N, Lundin D, Hofer A, Sjöberg BM, Logan DT. 2016a. Structural mode of allosteric activity regulation in a ribonucleotide reductase with duplicated ATP cones. Structure 24:906–917.

Johansson R, Jonna VR, Kumar R, Nayeri N, Lundin D, Sjöberg BM, Hofer A, Logan DT. 2016b. Structural Mechanism of Allosteric Activity Regulation in a Ribonucleotide Reductase with Double ATP Cones. Structure 24:906–917.

Jonna VR, Crona M, Rofougaran R, Lundin D, Johansson S, Brännström K, Sjöberg B-M, Hofer A. 2015. Diversity in Overall Activity Regulation of Ribonucleotide Reductase. J Biol Chem 290:17339– 17348.

Jordan A, Pontis E, Atta M, Krook M, Gibert I, Barbé J, Reichard P. 1994. A second class I ribonucleotide reductase in Enterobacteriaceae: characterization of the Salmonella typhimurium enzyme. Proc Natl Acad Sci U S A 91:12892–12896.

Jumper J, Evans R, Pritzel A, Green T, Figurnov M, Ronneberger O, Tunyasuvunakool K, Bates R, Žídek A, Potapenko A, Bridgland A, Meyer C, Kohl SAA, Ballard AJ, Cowie A, Romera-Paredes B, Nikolov S, Jain R, Adler J, Back T, Petersen S, Reiman D, Clancy E, Zielinski M, Steinegger M, Pacholska M, Berghammer T, Bodenstein S, Silver D, Vinyals O, Senior AW, Kavukcuoglu K, Kohli P, Hassabis D. 2021. Highly accurate protein structure prediction with AlphaFold. Nature 596:583–589.

Kalyaanamoorthy S, Minh BQ, Wong TKF, von Haeseler A, Jermiin LS. 2017. ModelFinder: fast model selection for accurate phylogenetic estimates. Nat Methods 14:587–589.

Kang G, Taguchi AT, Stubbe JA, Drennan CL. 2020. Structure of a trapped radical transfer pathway within a ribonucleotide reductase holocomplex. Science 368:424–427.

Larsson KM, Andersson J, Sjöberg BM, Nordlund P, Logan DT. 2001. Structural basis for allosteric substrate specificity regulation in anaerobic ribonucleotide reductases. Structure 9:739–750.

Larsson K-M, Jordan A, Eliasson R, Reichard P, Logan DT, Nordlund P. 2004. Structural mechanism of allosteric substrate specificity regulation in a ribonucleotide reductase. Nat Struct Mol Biol 11:1142–1149.

Larsson K-M, Logan DT, Nordlund P. 2010. Structural basis for adenosylcobalamin activation in AdoCbl-dependent ribonucleotide reductases. ACS Chem Biol 5:933–942.

Le SQ, Gascuel O. 2008. An improved general amino acid replacement matrix. Mol Biol Evol 25:1307– 1320.

Licht S, Gerfen GJ, Stubbe J. 1996. Thiyl radicals in ribonucleotide reductases. Science 271:477–481.

Lindell D, Jaffe JD, Coleman ML, Futschik ME, Axmann IM, Rector T, Kettler G, Sullivan MB, Steen R, Hess WR, Church GM, Chisholm SW. 2007. Genome-wide expression dynamics of a marine virus and host reveal features of co-evolution. Nature 449:83–86.

Lindell D, Jaffe JD, Johnson ZI, Church GM, Chisholm SW. 2005. Photosynthesis genes in marine viruses yield proteins during host infection. Nature 438:86–89.

Loderer C, Jonna VR, Crona M, Rozman Grinberg I, Sahlin M, Hofer A, Lundin D, Sjöberg B-M. 2017. A unique cysteine-rich zinc finger domain present in a majority of class II ribonucleotide reductases mediates catalytic turnover. J Biol Chem 292:19044–19054.

Logan DT, Andersson J, Sjöberg BM, Nordlund P. 1999. A glycyl radical site in the crystal structure of a class III ribonucleotide reductase. Science 283:1499–1504.

Logan DT, Mulliez E, Larsson K-M, Bodevin S, Atta M, Garnaud PE, Sjoberg B-M, Fontecave M. 2003. A metal-binding site in the catalytic subunit of anaerobic ribonucleotide reductase. Proc Natl Acad Sci U S A 100:3826–3831.

Lundin D, Berggren G, Logan D, Sjöberg B-M. 2015. The origin and evolution of ribonucleotide reduction. Life 5:604–636.

Lundin D, Torrents E, Poole AM, Sjöberg B-M. 2009. RNRdb, a curated database of the universal enzyme family ribonucleotide reductase, reveals a high level of misannotation in sequences deposited to Genbank. BMC Genomics 10:589.

Mao SS, Holler TP, Yu GX, Bollinger JM Jr, Booker S, Johnston MI, Stubbe J. 1992. A model for the role of multiple cysteine residues involved in ribonucleotide reduction: amazing and still confusing. Biochemistry 31:9733–9743.

Martínez-Carranza M, Jonna VR, Lundin D, Sahlin M, Carlson LA, Jemal N, Högbom M, Sjöberg BM, Stenmark P, Hofer A. 2020. A ribonucleotide reductase from clostridium botulinum reveals distinct evolutionary pathways to regulation via the overall activity site. J Biol Chem 295:15576–15587.

Meisburger SP, Thomas WC, Watkins MB, Ando N. 2017. X-ray Scattering Studies of Protein Structural Dynamics. Chemical Reviews. doi:10.1021/acs.chemrev.6b00790

Mulliez E, Ollagnier S, Fontecave M, Eliasson R, Reichard P. 1995. Formate is the hydrogen donor for the anaerobic ribonucleotide reductase from Escherichia coli. Proc Natl Acad Sci U S A 92:8759–8762.

Nakamura Y, Kaneko T, Sato S, Mimuro M, Miyashita H, Tsuchiya T, Sasamoto S, Watanabe A, Kawashima K, Kishida Y, Others. 2003. Complete genome structure of Gloeobacter violaceus PCC 7421, a cyanobacterium that lacks thylakoids. DNA Res 10:137–145.

Parker MJ. 2017. Discovery and investigation of the novel overall activity allosteric regulation of the Bacillus subtilis class Ib ribonucleotide reductase. Massachusetts Institute of Technology.

Parker MJ, Maggiolo AO, Thomas WC, Kim A, Meisburger SP, Ando N, Boal AK, Stubbe J. 2018. An endogenous dAMP ligand in Bacillus subtilis class Ib RNR promotes assembly of a noncanonical dimer for regulation by dATP. Proceedings of the National Academy of Sciences 115:E4594– E4603.

Pollock DD, Zwickl DJ, McGuire JA, Hillis DM. 2002. Increased taxon sampling is advantageous for phylogenetic inference. Syst Biol 51:664–671.

Poole AM, Logan DT, Sjöberg BM. 2002. The evolution of the ribonucleotide reductases: Much ado about oxygen. J Mol Evol 55:180–196.

Rehling D, Scaletti ER, Rozman Grinberg I, Lundin D, Sahlin M, Hofer A, Sjöberg B-M, Stenmark P. 2022. Structural and Biochemical Investigation of Class I Ribonucleotide Reductase from the Hyperthermophile Aquifex aeolicus. Biochemistry 61:92–106.

Reichard P. 1997. The evolution of ribonucleotide reduction. Trends Biochem Sci 22:81–85.

Reichard P. 1993. From RNA to DNA, why so many ribonucleotide reductases? Science 260:1773–1777.

Reichard P. 1988. Interactions between deoxyribonucleotide and DNA synthesis. Annu Rev Biochem 57:349–374.

Riera J, Robb FT, Weiss R, Fontecave M. 1997. Ribonucleotide reductase in the archaeon Pyrococcus furiosus: A critical enzyme in the evolution of DNA genomes? Proc Natl Acad Sci U S A 94:475– 478.

Rives A, Meier J, Sercu T, Goyal S, Lin Z, Liu J, Guo D, Ott M, Zitnick CL, Ma J, Fergus R. 2021. Biological structure and function emerge from scaling unsupervised learning to 250 million protein sequences. Proc Natl Acad Sci U S A 118. doi:10.1073/pnas.2016239118

Rose HR, Maggiolo AO, McBride MJ, Palowitch GM, Pandelia M-E, Davis KM, Yennawar NH, Boal AK. 2019. Structures of Class Id Ribonucleotide Reductase Catalytic Subunits Reveal a Minimal Architecture for Deoxynucleotide Biosynthesis 58:49.

Roshick C, Iliffe-Lee ER, McClarty G. 2000. Cloning and characterization of ribonucleotide reductase from Chlamydia trachomatis. J Biol Chem 275:38111–38119.

Rozman Grinberg I, Lundin D, Hasan M, Crona M, Jonna VR, Loderer C, Sahlin M, Markova N, Borovok I, Berggren G, Hofer A, Logan DT, Sjöberg BM. 2018. Novel ATP-cone-driven allosteric regulation of ribonucleotide reductase via the radical-generating subunit. Elife 7. doi:10.7554/eLife.31529

Ruskoski TB, Boal AK. 2021. The periodic table of ribonucleotide reductases. J Biol Chem 297:101137.

Saw JHW, Schatz M, Brown MV, Kunkel DD, Foster JS, Shick H, Christensen S, Hou S, Wan X, Donachie SP. 2013. Cultivation and complete genome sequencing of Gloeobacter kilaueensis sp. nov., from a lava cave in Kīlauea Caldera, Hawai’i. PLoS One 8:e76376.

Schell E, Nouairia G, Steiner E, Weber N, Lundin D, Loderer C. 2021. Structural determinants and distribution of phosphate specificity in ribonucleotide reductases. J Biol Chem 297:101008.

Schirrmeister BE, de Vos JM, Antonelli A, Bagheri HC. 2013. Evolution of multicellularity coincided with increased diversification of cyanobacteria and the Great Oxidation Event. Proc Natl Acad Sci U S A 110:1791–1796.

Seyedsayamdost MR, Chan CTY, Mugnaini V, Stubbe J, Bennati M. 2007. PELDOR spectroscopy with DOPA-β2 and NH2Y-α2s: Distance measurements between residues involved in the radical propagation pathway of E. coli ribonucleotide reductase. J Am Chem Soc 129:15748–15749.

Shimodaira H. 2002. An approximately unbiased test of phylogenetic tree selection. Syst Biol 51:492–508.

Simmons MP, Müller KF, Webb CT. 2011. The deterministic effects of alignment bias in phylogenetic inference. Cladistics 27:402–416.

Sintchak MD, Arjara G, Kellogg BA, Stubbe JA, Drennan CL. 2002. The crystal structure of class II ribonucleotide reductase reveals how an allosterically regulated monomer mimics a dimer. Nat Struct Biol 9:293–300.

Spence MA, Mortimer MD, Buckle AM, Minh BQ, Jackson CJ. 2021. A Comprehensive Phylogenetic Analysis of the Serpin Superfamily. Mol Biol Evol 38:2915–2929.

Srinivas V, Lebrette H, Lundin D, Kutin Y, Sahlin M, Lerche M, Eirich J, Branca RMM, Cox N, Sjöberg B-M, Högbom M. 2018. Metal-free ribonucleotide reduction powered by a DOPA radical in Mycoplasma pathogens. Nature 563:416–420.

Stubbe J. 2000. Ribonucleotide reductases: The link between an RNA and a DNA world? Current Opinion in Structural Biology. doi:10.1016/S0959-440X(00)00153-6

Stubbe J, Ge J, Yee CS. 2001. The evolution of ribonucleotide reduction revisited. Trends Biochem Sci 26:93–99.

Sullivan MB, Coleman ML, Weigele P, Rohwer F, Chisholm SW. 2005. Three Prochlorococcus cyanophage genomes: signature features and ecological interpretations. PLoS Biol 3:e144.

Sullivan MB, Lindell D, Lee JA, Thompson LR, Bielawski JP, Chisholm SW. 2006. Prevalence and evolution of core photosystem II genes in marine cyanobacterial viruses and their hosts. PLoS Biol 4:e234.

Thomas WC, Brooks FP 3rd, Burnim AA, Bacik J-P, Stubbe J, Kaelber JT, Chen JZ, Ando N. 2019. Convergent allostery in ribonucleotide reductase. Nat Commun 10:2653.

Thompson LR, Zeng Q, Kelly L, Huang KH, Singer AU, Stubbe JA, Chisholm SW. 2011. Phage auxiliary metabolic genes and the redirection of cyanobacterial host carbon metabolism. Proc Natl Acad Sci U S A 108. doi:10.1073/pnas.1102164108

Torrents E. 2014. Ribonucleotide reductases: essential enzymes for bacterial life. Front Cell Infect Microbiol 4. doi:10.3389/fcimb.2014.00052

Torrents E, Aloy P, Gibert I, Rodríguez-Trelles F. 2002. Ribonucleotide reductases: divergent evolution of an ancient enzyme. J Mol Evol 55:138–152.

Torrents E, Westman MA, Sahlin M, Sjöberg BM. 2006. Ribonucleotide Reductase Modularity: ATYPICAL DUPLICATION OF THE ATP-CONE DOMAIN IN PSEUDOMONAS AERUGINOSA. J Biol Chem 281:25287–25296.

Tully BJ, Graham ED, Heidelberg JF. 2018. The reconstruction of 2,631 draft metagenome-assembled genomes from the global oceans. Sci Data 5:170203.

Uhlin U, Eklund H. 1996. The ten-stranded beta/alpha barrel in ribonucleotide reductase protein R1. J Mol Biol 262:358–369.

Wei Y. 2015. Mechanistic investigations of class III anaerobic ribonucleotide reductases. Massachusetts Institute of Technology.

Wei Y, Funk MA, Rosado LA, Baek J, Drennan CL, Stubbe J. 2014a. The class III ribonucleotide reductase from Neisseria bacilliformis can utilize thioredoxin as a reductant. Proc Natl Acad Sci U S A 111:E3756–65.

Wei Y, Mathies G, Yokoyama K, Chen J, Griffin RG, Stubbe J. 2014b. A chemically competent thiosulfuranyl radical on the Escherichia coli class III ribonucleotide reductase. J Am Chem Soc 136:9001– 9013.

Weiss MC, Sousa FL, Mrnjavac N, Neukirchen S, Roettger M, Nelson-Sathi S, Martin WF. 2016. The physiology and habitat of the last universal common ancestor. Nature Microbiology 1. doi:10.1038/nmicrobiol.2016.116

Weiss RA. 2017. Exchange of Genetic Sequences Between Viruses and Hosts. Curr Top Microbiol Immunol 407:1–29.

Whelan S, Goldman N. 2001. A general empirical model of protein evolution derived from multiple protein families using a maximum-likelihood approach. Mol Biol Evol 18:691–699.

Zimanyi CM, Chen PYT, Kang G, Funk MA, Drennan CL. 2016. Molecular basis for allosteric specificity regulation in class ia ribonucleotide reductase from Escherichia coli. Elife 5:e07141.

## References

Afonine PV, Poon BK, Read RJ, Sobolev OV, Terwilliger TC, Urzhumtsev A, Adams PD. 2018. Realspace refinement inPHENIXfor cryo-EM and crystallography. Acta Crystallogr D Struct Biol 74:531–544.

Alley EC, Khimulya G, Biswas S, AlQuraishi M, Church GM. 2019. Unified rational protein engineering with sequence-based deep representation learning. Nat Methods 16:1315–1322.

Ando N, Brignole EJ, Zimanyi CM, Funk MA, Yokoyama K, Asturias FJ, Stubbe J, Drennan CL. 2011. Structural interconversions modulate activity of Escherichia coli ribonucleotide reductase. Proc Natl Acad Sci U S A 108:21046–21051.

Ando N, Li H, Brignole EJ, Thompson S, McLaughlin MI, Page JE, Asturias FJ, Stubbe J, Drennan CL. 2016. Allosteric Inhibition of Human Ribonucleotide Reductase by dATP Entails the Stabilization of a Hexamer. Biochemistry 55:373–381.

Asarnow, D., Palovcak, E., & Cheng, Y. 2019. asarnow/pyem: UCSF pyem v0.5. doi:10.5281/zenodo.3576630

Bateman A, Martin MJ, Orchard S, Magrane M, Agivetova R, Ahmad S, Alpi E, Bowler-Barnett EH, Britto R, Bursteinas B, Bye-A-Jee H, Coetzee R, Cukura A, Silva AD, Denny P, Dogan T, Ebenezer TG, Fan J, Castro LG, Garmiri P, Georghiou G, Gonzales L, Hatton-Ellis E, Hussein A, Ignatchenko A, Insana G, Ishtiaq R, Jokinen P, Joshi V, Jyothi D, Lock A, Lopez R, Luciani A, Luo J, Lussi Y, MacDougall A, Madeira F, Mahmoudy M, Menchi M, Mishra A, Moulang K, Nightingale A, Oliveira CS, Pundir S, Qi G, Raj S, Rice D, Lopez MR, Saidi R, Sampson J, Sawford T, Speretta E, Turner E, Tyagi N, Vasudev P, Volynkin V, Warner K, Watkins X, Zaru R, Zellner H, Bridge A, Poux S, Redaschi N, Aimo L, Argoud-Puy G, Auchincloss A, Axelsen K, Bansal P, Baratin D, Blatter MC, Bolleman J, Boutet E, Breuza L, Casals-Casas C, de Castro E, Echioukh KC, Coudert E, Cuche B, Doche M, Dornevil D, Estreicher A, Famiglietti ML, Feuermann M, Gasteiger E, Gehant S, Gerritsen V, Gos A, Gruaz-Gumowski N, Hinz U, Hulo C, Hyka-Nouspikel N, Jungo F, Keller G, Kerhornou A, Lara V, Le Mercier P, Lieberherr D, Lombardot T, Martin X, Masson P, Morgat A, Neto TB, Paesano S, Pedruzzi I, Pilbout S, Pourcel L, Pozzato M, Pruess M, Rivoire C, Sigrist C, Sonesson K, Stutz A, Sundaram S, Tognolli M, Verbregue L, Wu CH, Arighi CN, Arminski L, Chen C, Chen Y, Garavelli JS, Huang H, Laiho K, McGarvey P, Natale DA, Ross K, Vinayaka CR, Wang Q, Wang Y, Yeh LS, Zhang J. 2021. UniProt: The universal protein knowledgebase in 2021. Nucleic Acids Res 49:D480–D489.

Blum M, Chang H-Y, Chuguransky S, Grego T, Kandasaamy S, Mitchell A, Nuka G, Paysan-Lafosse T, Qureshi M, Raj S, Richardson L, Salazar GA, Williams L, Bork P, Bridge A, Gough J, Haft DH, Letunic I, Marchler-Bauer A, Mi H, Natale DA, Necci M, Orengo CA, Pandurangan AP, Rivoire C, Sigrist CJA, Sillitoe I, Thanki N, Thomas PD, Tosatto SCE, Wu CH, Bateman A, Finn RD. 2021. The InterPro protein families and domains database: 20 years on. Nucleic Acids Res 49:D344–D354.

Eddy SR. 2011. Accelerated Profile HMM Searches. PLoS Comput Biol 7:e1002195.

Edgar RC. 2004. MUSCLE: a multiple sequence alignment method with reduced time and space complexity. BMC Bioinformatics 5:113.

El-Gebali S, Mistry J, Bateman A, Eddy SR, Luciani A, Potter SC, Qureshi M, Richardson LJ, Salazar GA, Smart A, Sonnhammer ELL, Hirsh L, Paladin L, Piovesan D, Tosatto SCE, Finn RD. 2019. The Pfam protein families database in 2019. Nucleic Acids Res 47:D427–D432.

Emsley P, Cowtan K. 2004. Coot: model-building tools for molecular graphics. Acta Crystallogr D Biol Crystallogr 60:2126–2132.

Finn RD, Clements J, Eddy SR. 2011. HMMER web server: interactive sequence similarity searching. Nucleic Acids Res 39:W29–37.

Fu L, Niu B, Zhu Z, Wu S, Li W. 2012. CD-HIT: accelerated for clustering the next-generation sequencing data. Bioinformatics 28:3150–3152.

Hansen S. 2000. Bayesian estimation of hyperparameters for indirect Fourier transformation in small-angle scattering. J Appl Crystallogr 33:1415–1421.

Hie B, Zhong ED, Berger B, Bryson B. 2021. Learning the language of viral evolution and escape. Science 371:284–288.

Hoang DT, Chernomor O, von Haeseler A, Minh BQ, Vinh LS. 2018. UFBoot2: Improving the Ultrafast Bootstrap Approximation. Mol Biol Evol 35:518–522.

Holm L. 2020. Using Dali for Protein Structure Comparison In: Gáspári Z, editor. Structural Bioinformatics: Methods and Protocols. New York, NY: Springer US. pp. 29–42.

Hopkins JB, Gillilan RE, Skou S. 2017. BioXTAS RAW: improvements to a free open-source program for small-angle X-ray scattering data reduction and analysis. J Appl Crystallogr 50:1545–1553.

Katoh K, Rozewicki J, Yamada KD. 2017. MAFFT online service: multiple sequence alignment, interactive sequence choice and visualization. Brief Bioinform 20:1160–1166.

Konagurthu AS, Reboul CF, Schmidberger JW, Irving JA, Lesk AM, Stuckey PJ, Whisstock JC, Buckle AM. 2010. MUSTANG-MR structural sieving server: applications in protein structural analysis and crystallography. PLoS One 5:e10048.

Konarev PV, Volkov VV, Sokolova AV, Koch MHJ, Svergun DI. 2003. PRIMUS: a Windows PC-based system for small-angle scattering data analysis. J Appl Crystallogr 36:1277–1282.

Landau M, Mayrose I, Rosenberg Y, Glaser F, Martz E, Pupko T, Ben-Tal N. 2005. ConSurf 2005: the projection of evolutionary conservation scores of residues on protein structures. Nucleic Acids Res 33. doi:10.1093/nar/gki370

Liebschner D, Afonine PV, Moriarty NW, Poon BK, Chen VB, Adams PD. 2021. CERES: a cryo-EM rerefinement system for continuous improvement of deposited models. Acta Crystallogr D Struct Biol 77:48–61.

Manalastas-Cantos K, Konarev PV, Hajizadeh NR, Kikhney AG, Petoukhov MV, Molodenskiy DS, Panjkovich A, Mertens HDT, Gruzinov A, Borges C, Jeffries CM, Svergun DI, Franke D. 2021. ATSAS 3.0: expanded functionality and new tools for small-angle scattering data analysis. J Appl Crystallogr 54:343–355.

McInnes, L., Healy, J., Saul, N., & Großberger, L. 2018. UMAP: Uniform Manifold Approximation and Projection. Journal of Open Source Software 3:861.

Meisburger SP, Taylor AB, Khan CA, Zhang S, Fitzpatrick PF, Ando N. 2016. Domain Movements upon Activation of Phenylalanine Hydroxylase Characterized by Crystallography and Chromatography-Coupled Small-Angle X-ray Scattering. J Am Chem Soc 138:6506–6516.

Meisburger SP, Xu D, Ando N. 2021. REGALS: a general method to deconvolve X-ray scattering data from evolving mixtures. IUCrJ 8:225–237.

Minh BQ, Schmidt HA, Chernomor O, Schrempf D, Woodhams MD, von Haeseler A, Lanfear R. 2020. IQ-TREE 2: New Models and Efficient Methods for Phylogenetic Inference in the Genomic Era. Mol Biol Evol 37:1530–1534.

Nguyen N-PD, Mirarab S, Kumar K, Warnow T. 2015. Ultra-large alignments using phylogeny-aware profiles. Genome Biol 16:124.

Parker MJ. 2017. Discovery and Investigation of the Novel Overall Activity Allosteric Regulation of the Bacillus subtilis Class Ib Ribonucleotide Reductase (Doctor of Philosophy in Biological Chemistry). Massachusetts Institute of Technology.

Punjani A, Rubinstein JL, Fleet DJ, Brubaker MA. 2017. cryoSPARC: algorithms for rapid unsupervised cryo-EM structure determination. Nat Methods 14:290–296.

Punjani A, Zhang H, Fleet DJ. 2020. Non-uniform refinement: adaptive regularization improves singleparticle cryo-EM reconstruction. Nat Methods 17:1214–1221.

Rives A, Meier J, Sercu T, Goyal S, Lin Z, Liu J, Guo D, Ott M, Zitnick CL, Ma J, Fergus R. 2021. Biological structure and function emerge from scaling unsupervised learning to 250 million protein sequences. Proc Natl Acad Sci U S A 118:e2016239118.

Schneidman-Duhovny D, Hammel M, Sali A. 2010. FoXS: a web server for rapid computation and fitting of SAXS profiles. Nucleic Acids Res 38:W540–4.

Shannon P, Markiel A, Ozier O, Baliga NS, Wang JT, Ramage D, Amin N, Schwikowski B, Ideker T. 2003. Cytoscape: A Software Environment for Integrated Models. Genome Res 13:2498–2504.

Sievers F, Wilm A, Dineen D, Gibson TJ, Karplus K, Li W, Lopez R, McWilliam H, Remmert M, Söding J, Thompson JD, Higgins DG. 2011. Fast, scalable generation of high-quality protein multiple sequence alignments using Clustal Omega. Mol Syst Biol 7:539.

Skou S, Gillilan RE, Ando N. 2014. Synchrotron-based small-angle X-ray scattering of proteins in solution. Nat Protoc 9:1727–1739.

Spence MA, Mortimer MD, Buckle AM, Minh BQ, Jackson CJ. 2021. A comprehensive phylogenetic analysis of the serpin superfamily. Mol Biol Evol. doi:10.1093/molbev/msab081

Svergun D, Barberato C, Koch MHJ. 1995. CRYSOL – a Program to Evaluate X-ray Solution Scattering of Biological Macromolecules from Atomic Coordinates. J Appl Crystallogr 28:768–773.

Thomas WC, Brooks FP, Burnim AA, Bacik JP, Stubbe JA, Kaelber JT, Chen JZ, Ando N. 2019. Convergent allostery in ribonucleotide reductase. Nat Commun 10:2653.

Weinkam P, Pons J, Sali A. 2012. Structure-based model of allostery predicts coupling between distant sites. Proc Natl Acad Sci U S A 109:4875–4880.

Zallot R, Oberg N, Gerlt JA. 2019. The EFI Web Resource for Genomic Enzymology Tools: Leveraging Protein, Genome, and Metagenome Databases to Discover Novel Enzymes and Metabolic Pathways. Biochemistry 58:4169–4182.

Zheng SQ, Palovcak E, Armache J-P, Verba KA, Cheng Y, Agard DA. 2017. MotionCor2: anisotropic correction of beam-induced motion for improved cryo-electron microscopy. Nat Methods 14:331– 332.

Zivanov J, Nakane T, Forsberg BO, Kimanius D, Hagen WJ, Lindahl E, Scheres SH. 2018. New tools for automated high-resolution cryo-EM structure determination in RELION-3. Elife 7. doi:10.7554/eLife.42166

Zivanov J, Nakane T, Scheres SHW. 2020. Estimation of high-order aberrations and anisotropic magnification from cryo-EM data sets in RELION-3.1. IUCrJ 7:253–267.

Zivanov J, Nakane T, Scheres SHW. 2019. A Bayesian approach to beam-induced motion correction in cryo-EM single-particle analysis. IUCrJ 6:5–17.

